# Pathway-based similarity measurement to quantify transcriptomics similarity between human tissues and preclinical models

**DOI:** 10.1101/2024.04.13.589385

**Authors:** Paarth Parekh, Jason Sherfey, Begum Alaybeyoglu, Murat Cirit

## Abstract

Accurate clinical translation of preclinical research remains challenging, primarily due to species-specific differences and disease and patient heterogeneity. An important recent advancement has been development of microphysiological systems that consist of multiple human cell types that recapitulate key characteristics of their respective human systems, allowing essential physiologic processes to be accurately assessed during drug development. However, an unmet need remains regarding a quantitative method to evaluate the similarity between diverse sample types for various contexts of use (CoU)-specific pathways. To address this gap, this study describes the development of pathway-based similarity measurement (PBSM), which leverages RNA-seq data and pathway-based information to assess the human relevance of preclinical models for specific CoU. PBSM offers a quantitative method to compare the transcriptomic similarity of preclinical models to human tissues, shown here as proof of concept for liver and cardiac tissues, enabling improved model selection and validation. Thus, PBSM can successfully support CoU selection for preclinical models, assess the impact of different gene sets on similarity calculations, and differentiate among various *in vitro* and *in vivo* models. PBSM has potential to reduce the translational gap in drug development by allowing quantitative evaluation of the similarity of preclinical models to human tissues, facilitating model selection, and improving understanding of context-specific applications. PBSM can serve as a foundation for enhancing the physiological relevance of *in vitro* models and supporting the development of more effective therapeutic interventions.

## Introduction

The translational gap between preclinical and clinical research remains a critical challenge in the development of novel therapeutic interventions. While preclinical studies frequently yield encouraging results in cellular and animal models, these findings may not always translate into clinical efficacy and safety in human subjects. The existence of species-specific differences and heterogeneity in disease among individual patients and heterogeneity across modeling systems and patients complicate the successful transition of preclinical discoveries into clinically effective treatments.

To address this challenge, there has been a concerted effort over the last decade to develop sophisticated human-based complex *in vitro* models known as microphysiological systems (MPS). These models offer a potential avenue for bridging the translational gap, particularly in well-defined contexts of use (CoUs). Significantly, the 2022 FDA Modernization Act recognizes human-based complex *in vitro* models as New Approach Methods (NAMs) that are poised to improve on the performance of current preclinical *in vitro* and animal models. An integral part of this development and implementation of new research tools or methods in drug development lies in their qualification for a specific CoU and validation for precision, accuracy, and reproducibility. A crucial aspect of the qualification and validation of NAMs is demonstration of human physiological relevance. Current methodologies for evaluating the human relevance of preclinical models rely on comparing a limited set of qualitative and quantitative physiological metrics. For instance, in the case of preclinical liver models, morphological metrics encompass aspects such tissue polarization, cell-cell junction maturity, and organellar content, while functional evaluation includes analysis of the secretome (albumin, urea, and acute phase protein secretion), functional assays (drug metabolizing activity assays), and responsiveness to injury (ALT increase). While these metrics serve both as measures of physiological relevance and quality control in experiments, there is a significant unmet need for a systematic and quantitative approach to assess the human relevance of preclinical models. Transcriptomics provide a broader scope of relevant biology than a few specific morphological and functional endpoints and they assess the genetic and epigenetic machinery that defines how a cell responds to usual physiological demands or injury. In gene expression profiling, RNA-seq methodology is particularly useful for characterizing genome functionality, organellar functions, and providing insights into developmental processes and diseases[1]. While RNA-seq is commonly employed to compare differences in gene expression profiles[2], it can also serve as a valuable tool for estimating the similarities among transcriptomic signatures.

Prior efforts to assess similarity using RNA-seq data have led to the development of tools like CellNet[3], TROM[4], and LiGEP[5]. However, these tools have several limitations, including a lack of flexibility and customizability, and are often tailored for specific tissue types. Furthermore, they fall short in providing a quantitative means to evaluate the similarity between diverse sample types for various CoU-specific pathways.

In response to these challenges, we introduce the Pathway-Based Similarity Measurement (PBSM), depicted in Figure 1. PBSM represents a pathway-based approach designed to quantitatively measure transcriptomic similarity between distinct biological samples. Leveraging various pathways, PBSM can comprehensively assess the similarity between samples by performing pairwise comparisons, ultimately generating a quantifiable similarity score. Moreover, PBSM holds the potential to validate a CoU by comparing gene expression similarities with human tissues, focusing on pathways vital for a given CoU. This study thus presents a novel and versatile approach for addressing the unmet need for quantitative assessment of human relevance of preclinical models.

**Figure 1:**
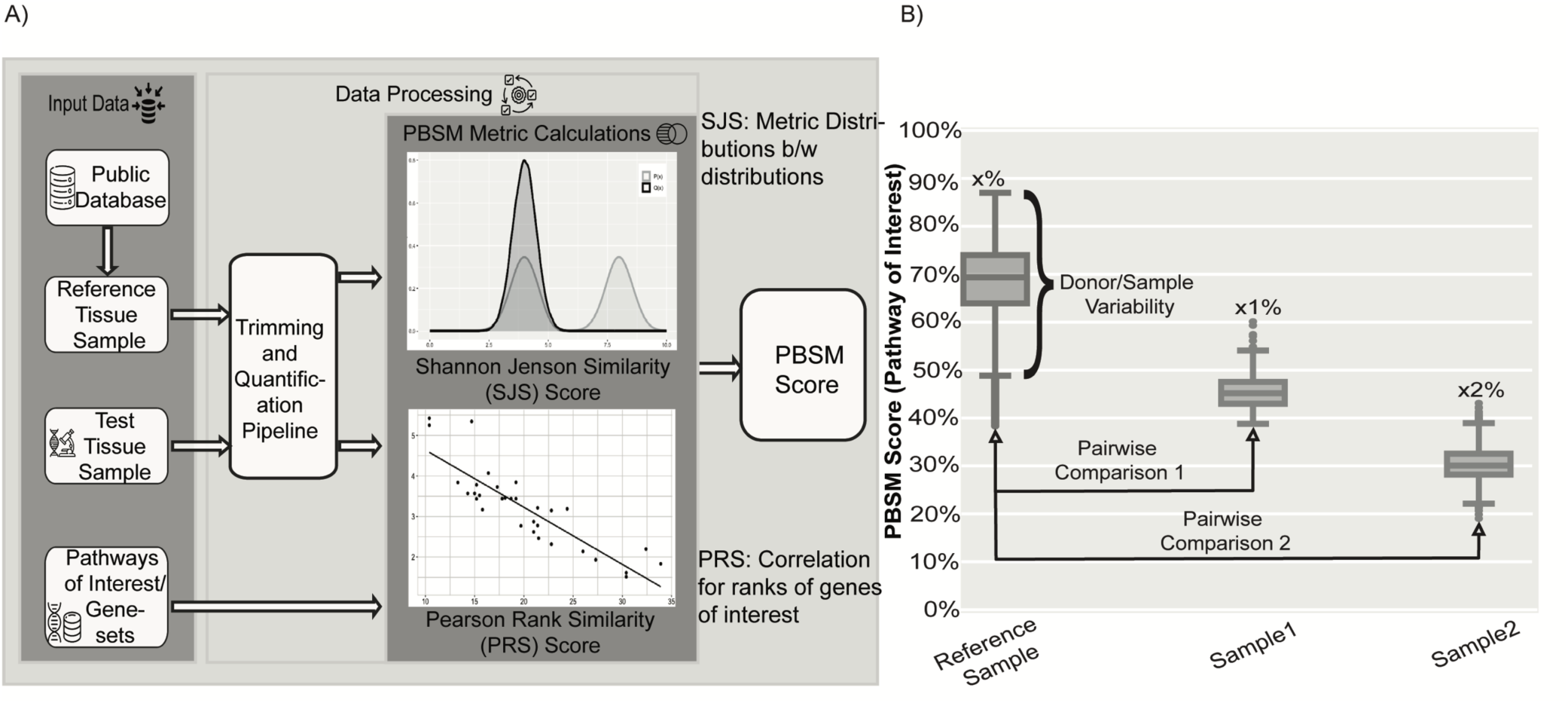
Outline for the Pathway-Based Similarity Metric (PBSM). A) Illustrates the pipeline and the data needed by PBSM to calculate the similarity between the RTS and TTS data using pathways and genes of interest. B) Illustrates an example plot indicating how to interpret the results generated throughout the article.

## Methods

### Data Sources and Processing

The Genotype-Tissue Expression (GTEx) database by Broad Institute of Science was used to obtain the RNA-seq datasets for human tissue samples. Detailed analysis for the GTEX is provided in the supplementary information.

Publicly available RNA-seq datasets for various preclinical models used in PBSM were obtained from Gene Expression Omnibus (GEO). These RNA-seq datasets were processed to obtain normalized reads and were then benchmarked with the corresponding GTEx tissue samples to generate median similarity scores. Detailed data processing steps are provided in the supplementary material.

Transcript Per Million (TPMs) was used to calculate the PBSM score for the pairwise gene expression comparison. This minimizes the bias introduced by differences in library sizes and gene lengths. While TPMs do not account for the RNA library composition, the ability to add more samples to compare the expression between samples over time and equal library size plays a more important role than the need to account for library composition[6].

### PBSM methodology

PBSM requires pathway information to provide the basis for the similarity calculations. This pathway information in the form of gene sets can be obtained through various databases like KEGG[7], Reactome[8], and MsigDb[9]. The input for the PBSM could be one or a list of combined gene sets from various pathways. Custom gene sets can also be generated for pairwise comparison of the two RNA-seq transcriptomics data sets.

PBSM takes the normalized gene counts, and the pathway/pathways information (PI) as input to calculate the median similarity score between the samples. PBSM measures the similarity score between two biological samples, such as human tissues and human MPSs, the reference tissue sample (RTS), and the testing tissue sample (TTS) data. PBSM consists of two metrics, the Shannon Jensen Similarity (SJS) and the Pearson Ranked Similarity (PRS), giving equal weight to both to obtain the PBSM score.

PBSM is scored based on the sample-to-sample comparison between the reference tissue samples and the testing tissue samples, taking the median similarity scores for genes expressed in a particular pathway for the two sets of samples.

Probability distribution and rank distribution of the gene sets:

The two TPM datasets, RTS and TTS, were converted into their subsequent probability for each gene based on the expression levels according to the Tissue probability equation (*T_pro_*), and the subset of genes present in the PI was selected from both the datasets for further analysis. Further ranks were calculated for the genes in the PI from the RNA-seq transcriptomics data, making it robust to outliers for further computation.

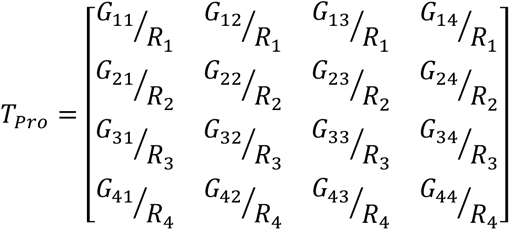

Shannon Jensen Similarity (SJS):

SJS is a widely utilized metric for quantifying the similarity between probability distributions, particularly in the context of information retrieval[10] and computational biology[11]. It is obtained from the Jensen-Shannon Divergence (JSD), measuring the distance between two probability distributions. JSD combines the Kullback-Leibler Divergence (KLD) and symmetric KLD to obtain a symmetric measure of similarity[12]. JSD is bounded by 1 when used with base 2 logarithm[12]. Previously, JSD has been used in a variety of bioinformatics[13] and genomics research[14][15]. We calculated the JSD between the two probability distributions of the RTS and TTS genes as follows:

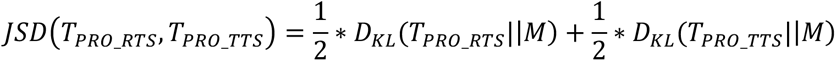

where

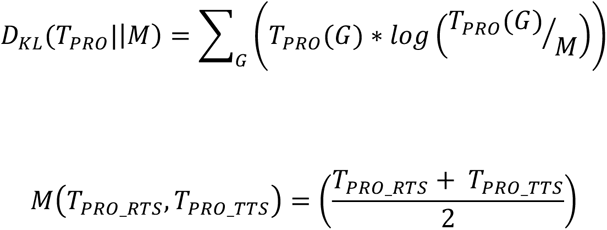

Shannon Jensen Distance (SJD) is the square root of JSD. This SJD is also bounded by 1, and we could thus obtain the SJS by subtracting 1 from the SJD and getting the similarity between two gene sets in terms of their probability distributions.

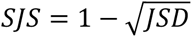

Pearson Rank Correlation (PRC):

PRC is a non-parametric measure of the monotonic correlation relationship between two variables. While the Shannon-Jenson similarity measured how similar the two data sets or objects are, correlation with gene expression data helps to measure the statistical relationship between the pairwise gene sets [16].

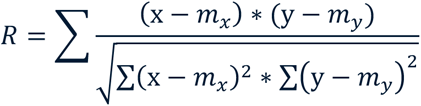

PRC is a form of the Pearson correlation, using the rank data to establish a correlation with the value ranging from 0 to 1. The value 0 indicates a negative correlation while a value of 1 indicates a positive correlation.

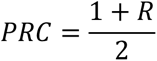

Taking the ranked subset of genes over the actual values makes it robust to handling outliers in the data and making it possible to conduct a non-parametric analysis while making no assumptions about the data [16].

## Results

### Evaluation of PBSM using the GTEx datasets reveals the high specificity and sensitivity of median similarity scores

Assessment of the bioinformatics workflow is crucial to ensure the accuracy and sensitivity of the obtained similarity scores. This entails verifying that the PBSM measures the intended parameters and produces results that are both reliable and reproducible. It should also possess the sensitivity required to detect meaningful variations among different gene sets and distinctions between diverse sample types. For this evaluation, publicly available GTEx tissue-specific transcriptomics datasets were used. Median similarity scores were computed through pairwise comparisons of RNA-seq datasets from various tissues, including brain, colon, heart, kidney, liver, muscle, and small intestine samples. The foundation for these calculations was established using sets of genes identified by the Human Protein Atlas (HPA)[17] as either highly organ-specific (HS) or highly expressed (HE) in individual organs.

Figure 2a and 2b present the median similarity scores obtained for different human organ samples based on HS and HE gene sets, respectively. Similarity scores were computed by comparing organ samples, with the reference organ designated as RTS (y-axis) and the compared organs as TTS (x-axis). In Figure 2a and 2b, the highest median similarity scores are observed along the diagonals. These diagonals represent cases where the RTS and TTS are the same, and the similarity score captures the differences among tissue donors within the same organ.

**Figure 2:**
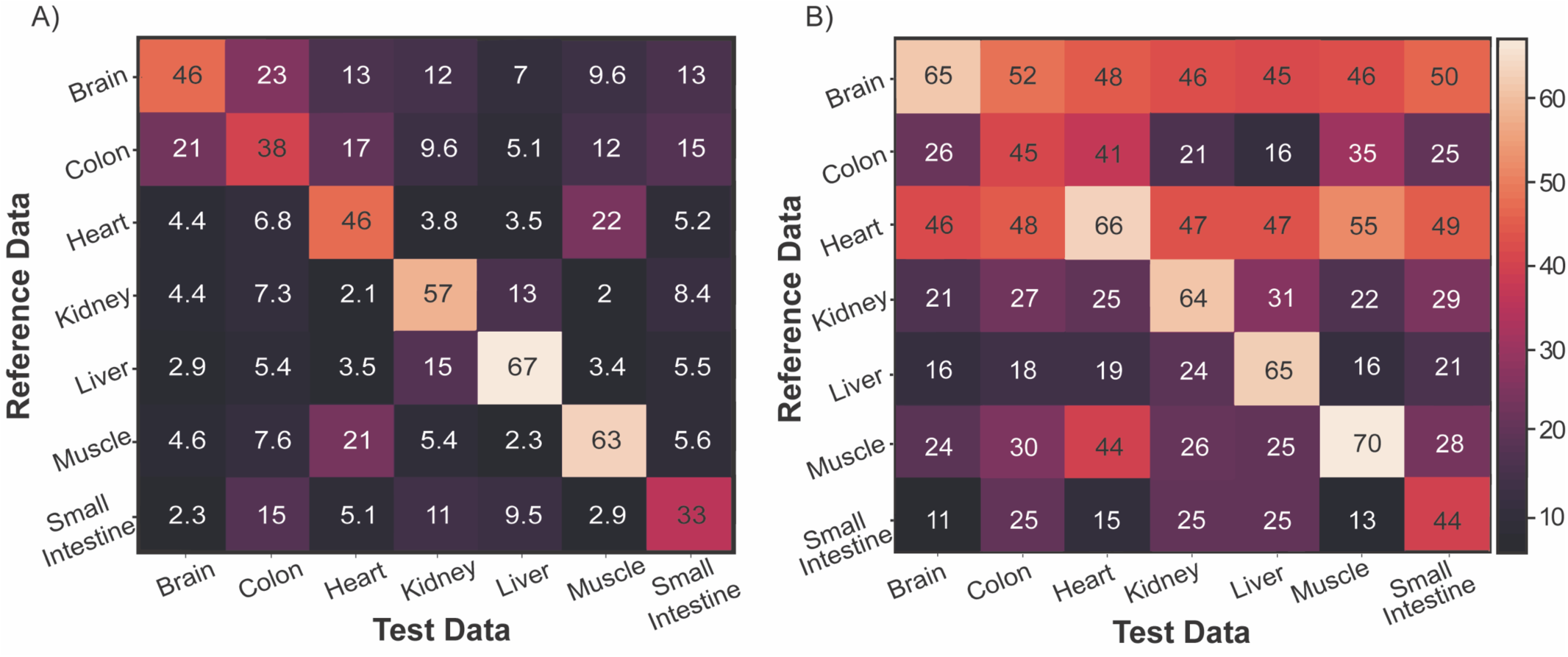
Assessment of tissue-specificity using the PBSM algorithm for the organ-specific & highly expressed gene sets. A) Median similarity scores that are obtained using the organ-specific gene set across the different RTS and TTS tissue samples. B) Median similarity scores that were obtained for the highly expressed gene set across the different RTS and TTS tissue samples.

The distinctions in median similarity scores between the two sub-figures in Figure 2 elucidate the sensitivity and specificity of PBSM. Figure 2a illustrates the similarity scores between organs based on HS genes, revealing the specificity of scores. Low scores between any given reference organ and different organs (see low values off the diagonal) confirm that the PBSM, using highly specific genes, accurately identifies dissimilar organs. Sensitivity is evident in the relatively large values of similarity between any given organ and itself (shown by large values on the diagonals) for both HS and HE genes in Figure 2a and 2b, respectively. Notably, the specificity of highly-expressed genes is lower (see large values off-diagonal in Figure 2b), as highly expressed genes, are not as organ-specific as highly-specific genes. Figure 2a further emphasizes sensitivity by featuring higher median similarity scores along the diagonals but displaying an overall lower median similarity score for other organs compared to HE genes in Figure 2b. This underscores PBSM’s sensitivity and the effects of using different gene sets to compare samples.

In summary, PBSM successfully estimated differences and similarities in RNA-seq organ data by employing organ-specific and highly expressed gene sets. It effectively demonstrated both the sensitivity and specificity of the tool and underscored the importance of the selected pathways/gene set used when comparing samples.

### Effect of different ADME gene sets on similarity calculations among organ datasets

To assess the effect of gene sets on PBSM-based similarity scores, we employed the absorption, distribution, metabolism, and excretion (ADME) pathway[18] as a case study. PBSM-based similarity scores for the Extended ADME pathway[19][Table S1] were computed using the RNA-seq datasets[Figure S2]. Using the GSEA overlap feature, various pathways were obtained from databases related to this extended ADME gene set[19][Table S1] [Figure S3], including from the KEGG[7][Table S2] and Reactome[8][Table S3] databases. Figure 3a illustrates the median similarity scores resulting from pairwise comparisons between the Reference Human Tissue Liver samples and other organs presented on the x-axis; the various ADME pathway gene sets sourced from different databases are shown on the y-axis. Despite their diverse origins, these pathways share involvement in the ADME processes. PBSM consistently reveals a similar pattern across these different gene sets: the liver tissue sample exhibits the highest similarity to the small intestine and kidney tissue samples for this process, as observed across the four overlapping ADME-related gene sets. Figure 3b shows the disparities and commonalities between the ADME pathway gene sets obtained from the various databases.

**Figure 3:**
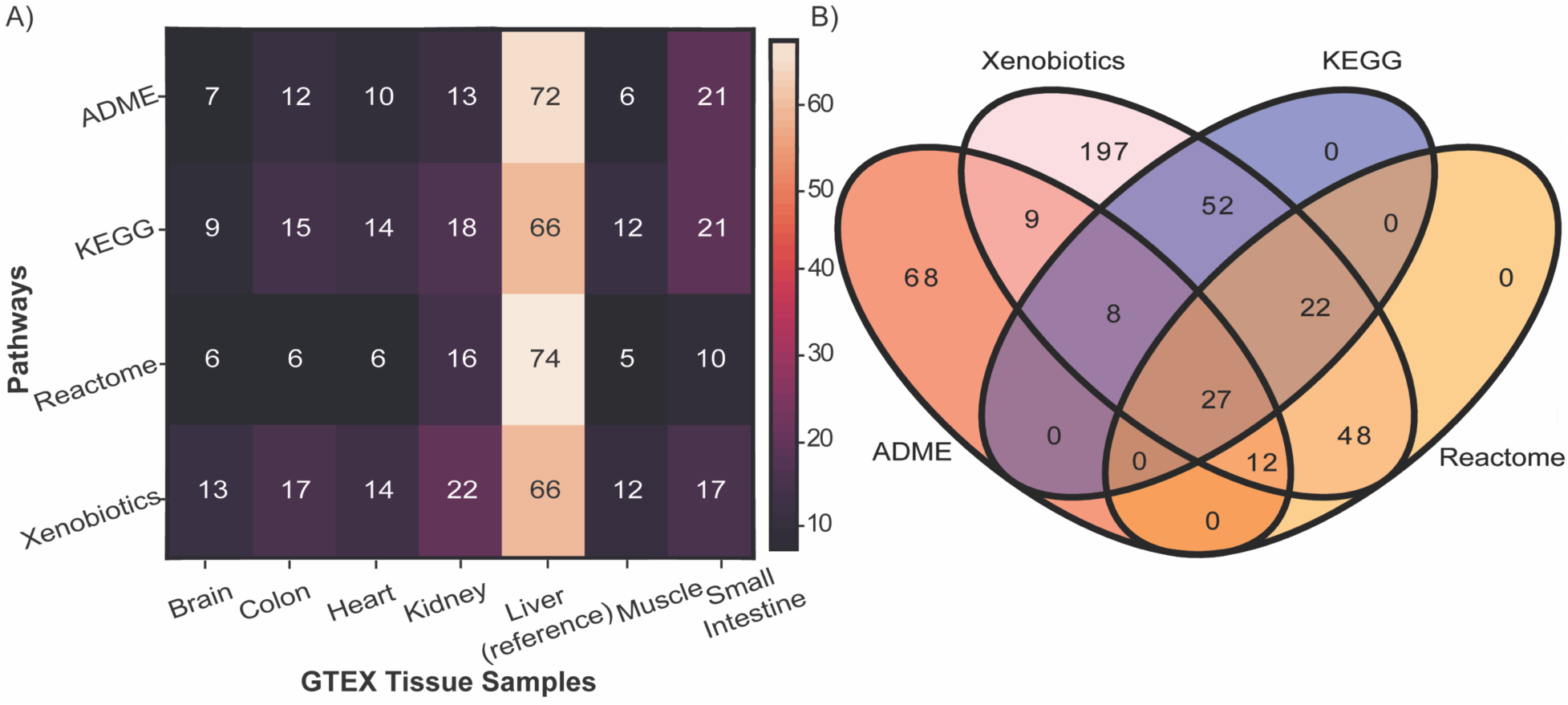
Assessment of the effect of ADME gene sets obtained from various databases on the PBSM-based similarity score. A) Heatmap for median similarity scores for the different ADME pathway databases on the y-axis indicating the differences in the similarity scores between the different organ types on the x-axis. B) Venn diagram showing the gene distribution for ADME pathways from different databases.

This demonstrates that PBSM-based similarity scores exhibit a degree of resilience to the source of gene sets for the ADME pathway. Moreover, the differences in liver-to-liver similarity (as seen in the Reference Liver column) reflect the differences in donor variability when defining gene sets related to the ADME pathway through alternative means. Further, the pairwise calculations between samples ensure that the variability of all the RTS donor information is captured.

### Comparing similarity scores for various *in vitro* and *in vivo* preclinical models to human liver using the extended ADME gene-set

We leveraged RNA-seq data from multiple preclinical *in vitro* and *in vivo* models commonly used in ADME studies to compute PBSM-based similarity scores with reference human biopsies based on the ADME pathway. Pairwise similarity calculations were conducted between the reference 220 liver donor samples from GTEx and the *in vitro* data illustrated in Figure 4a, sourced from GSE140520[20]. The ADME pathway, as detailed above and defined by the Extended ADME gene set[19][Table S3], served as the basis for comparing the similarity scores across these diverse models.

**Figure 4:**
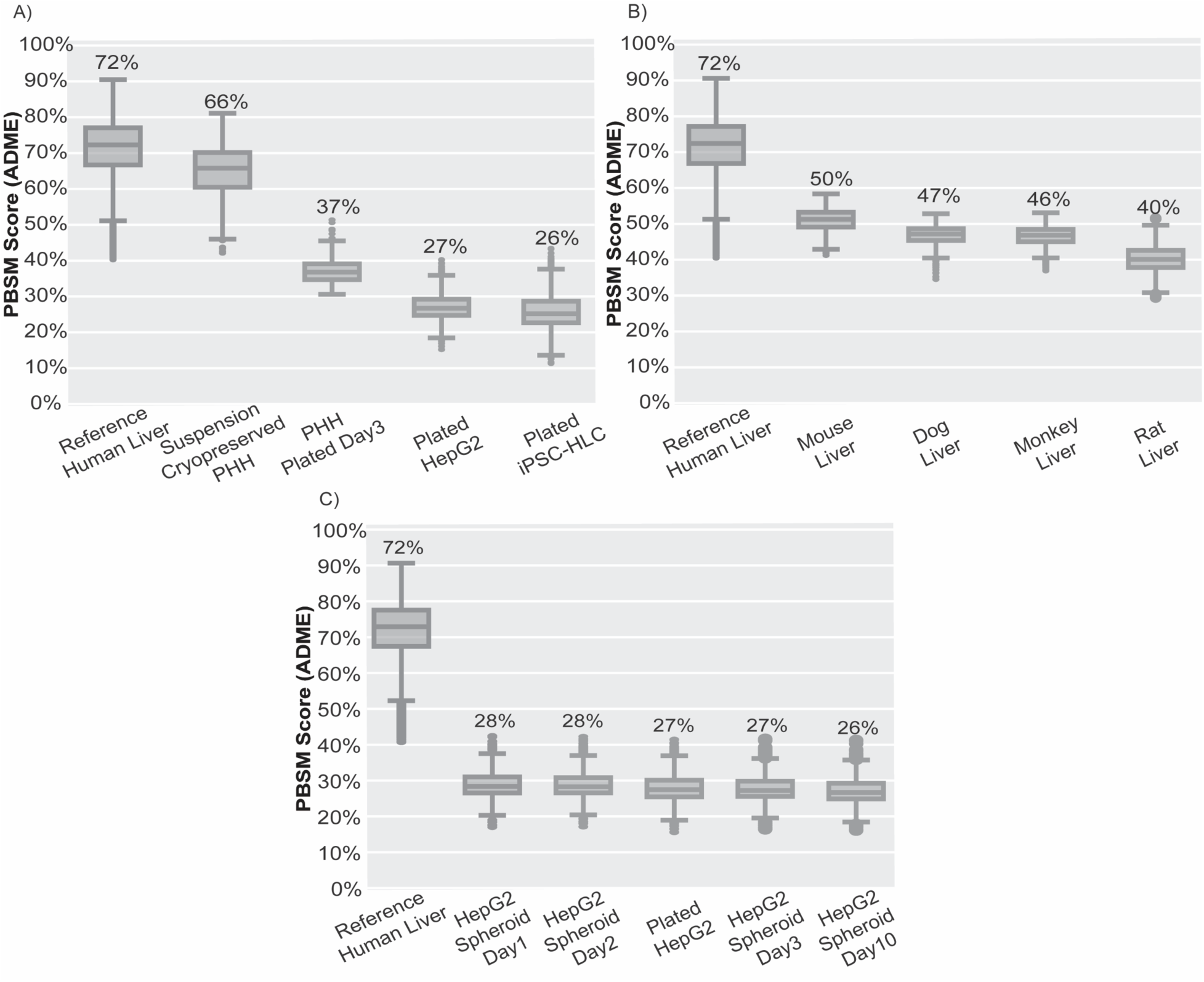
PBSM-based similarity scores for the various preclinical models when compared to the reference human liver biopsies using the extended ADME gene-set. A) Four in-vitro hepatic models are benchmarked against the reference human liver B) Four preclinical in-vivo models are benchmarked against the reference human liver. C) HepG2 model in 2D and 3D (spheroid) cultures were benchmarked against the reference human liver.

In Figure 4a, suspension PHH exhibited the highest median similarity score compared to the human liver reference. In contrast, plated PHH displayed similarity was approximately 50% lower after 3 days. Although the median similarity scores for the 3-day plated culture were lower than those of suspension PHH, they still maintained higher levels of similarity compared to other available *in vitro* models, such as plated HepG2 and iPSC-derived hepatocyte-like cells (iPSC-HLC).

Datasets for liver samples derived from various preclinical animal models employed in ADME studies were obtained from GEO. The *in vivo* datasets were aggregated from multiple sources, including mouse data from GSE222171[21], rat data from GSE55347[22], and dog and monkey data from GSE162142[23]. Figure 4b illustrates the similarity scores for liver datasets from four animal models (mouse, rat, dog, and monkey). Among these models, liver samples from mice exhibited the highest median similarity score to the reference human liver. Dog and monkey liver samples exhibit comparable levels of similarity, while the lowest scores were observed for rat liver samples.

Figure 4c explores the effects of culture type (2D vs. 3D HepG2) and duration in 3D cultures[24] using PBSM to calculate similarity scores to the reference human liver, employing the same extended ADME pathway. Pairwise comparisons to human liver were conducted for the 2D *in vitro* data from GSE140520[20] and the multiple-day 3D spheroid data of the HepG2 cell line from GSE213944[25]. The median similarity scores for both 2D and 3D HepG2 cultures over time were consistently low and independent of culture type and duration.

### Assessment of similarity scores of 2D and 3D iPSC-derived cardiomyocyte cultures using gene sets for cardiac-specific and drug-induced QT prolongation

In Figure 5a and 5b, PBSM scores were calculated to assess the maturation and long-term culture effects for the iPSC-derived cardiomyocytes (iPSC-CM), cultured in either 2D or 3D formats. Atrial and ventricle tissue reference samples sourced from the GTEx data were used for Figure 5a and 5b, respectively. Pairwise similarity scores were calculated between these reference samples and publicly available datasets for 2D and 3D iPSC-CM cultures obtained from GEO for maturation (GSE116574[26]) and long-term culture studies (GSE209997[27]).

**Figure 5:**
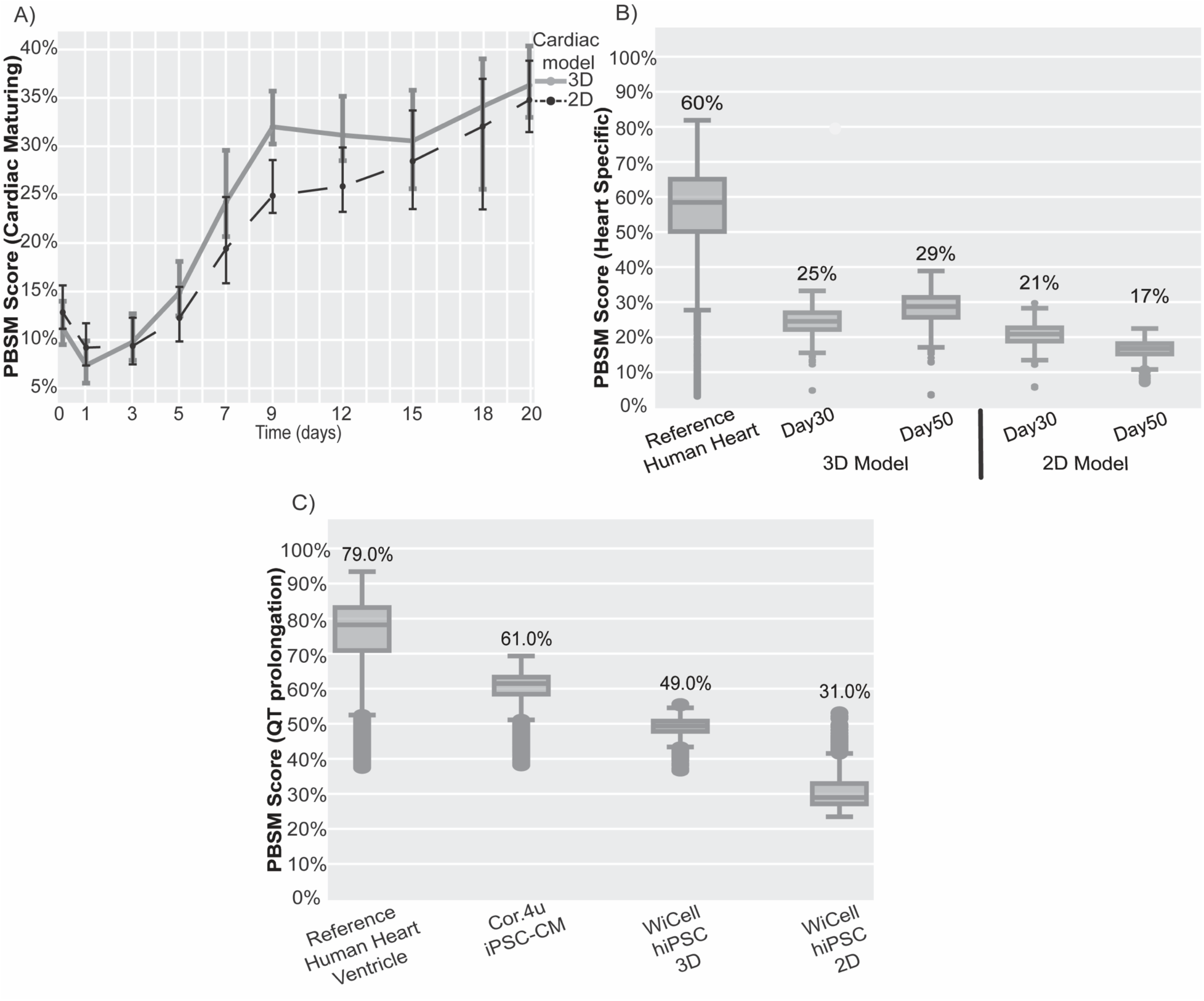
PBSM-based similarity scores estimated for the iPSC-derived cardiomyocytes A) Median similarity scores for both the 2D and 3D models during in-vitro maturation for 20 days using custom cardiac-related genes for the reference human atrial tissue samples. B) Median similarity scores showing the effects of the long-term culture for the 2D and 3D cardiomyocytes until 50 days for the reference human ventricle tissue Samples. C) Benchmarked three iPSC models to obtain median similarity scores compared to the reference samples to predict the ability for drug-induced QT-prolongation.

Figure 5a illustrates alterations in similarity scores during the iPSC-CM maturation period in both 2D and 3D cultures. The x-axis represents the maturation time points, while the y-axis represents the median similarity scores computed between the transcriptomics data from human atrial samples (reference human heart) and the iPSC-CM transcriptomics data. For these calculations, a custom gene set consisting of 254 genes [Table S4], encompassing genes involved in cardiac differentiation progression, CM functional and structural maturation, CM metabolism, WNT, TGF-β, and FGF signaling pathways, was employed.

Comparison between 2D vs 3D cultures during the maturation period revealed that similarity scores in both models increased approximately three-fold over time. Remarkably, the iPSC-CM 3D cultures achieved higher similarity scores at a faster rate than iPSC-CM 2D cultures. While both models reached similar levels of median similarity scores by day 20, the 3D model attained higher similarity levels by day 9 and maintained these elevated similarity scores.

Figure 5b investigates the impact of long-term culture on similarity scores for iPSC-CM 2D and 3D cultures. Pairwise comparisons were conducted between the reference ventricle datasets from GTEx and the long-term iPSC-CM cultures. In this case, the heart-specific gene set defined by the HPA[17] was employed to compute PBSM similarity scores. The results indicated that the long-term 3D culture study exhibits higher similarity scores than long-term 2D culture on days 30 and 50. Furthermore, 3D cultures showed a continued increase in similarity scores between days 30 and 50, while the scores for 2D cultures decreased significantly by day 50.

A pivotal aspect of the drug development process involves evaluating a drug’s potential to induce life-threatening cardiac arrhythmias, particularly QT prolongation. Previous studies have highlighted the promising correlation between Cor.4u-induced pluripotent stem cell cardiomyocytes (iPSC-CM) and clinical trial outcomes for drugs inducing QT prolongation [28]. Pairwise similarity scores were computed between the Human Reference Ventricle Cardiac Samples (RTS) and the publicly available datasets from GEO (TTS), encompassing Cor.4u iPSC-CM datasets (GSE114686[29])and the WiCell human-induced pluripotent stem cells (hiPSC) differentiated toward the cardiovascular lineage dataset for both 2D and 3D models (GSE209997[27]). In Figure 5c, Cor.4u iPSC-CM exhibits higher similarity than the 2D and 3D models for WiCell hiPSC to the human reference ventricle samples. The multiple cardiac ion channel pathways and genes associated with prolongation of QT interval [Table S5] were used for this comparison.

Thus, we observed strong correlation between PBSM-derived similarity scores and the capacity of Cor.4u iPSC-CM cells in identifying drugs that induce QT prolongation. Additionally, it highlights the potential of PBSM in discerning differences between various *in vitro* iPSC-CM models for a specific CoU.

### Assessment of PBSM-based similarity scores to distinguish preclinical models in disease contexts

One of the major translational challenges is the recapitulation of characteristics of human diseases in preclinical models[30]. Here, we investigated three preclinical animal models frequently used in non-alcoholic fatty liver (NAFL) and non-alcoholic steatohepatitis (NASH) research and estimated PBSM-based similarity scores through pairwise comparisons with datasets of human liver samples collected from NAFL or NASH patients, respectively. The reference datasets for human NAFL and NASH samples were sourced from GSE167523[31]. The datasets for preclinical animal models were obtained from GEO, encompassing the mouse high-fat model (GSE222171[20]), mice injected with CCl4 (GSE207855[32]), and the rat high-fat model (GSE192425[33]). For our analysis, three distinct pathways corresponding to gene sets were used: liver-specific genes sourced from HPA[17], the KEGG NAFLD pathway[Table S6], and a custom gene set involving genes associated with fibrosis [Table S7].

In Figure 6a, the use of liver-specific genes from the HPA resulted in consistently low and comparable similarity scores for each preclinical animal model when benchmarked against the human NAFL reference. In contrast, Figure 6b and 6c, used disease-related pathways and gene sets. Figure 6b revealed that the mouse high-fat model exhibited the highest median similarity scores compared to the datasets from NAFL patient samples using NAFLD gene set. Conversely, Figure 6c, utilizing the fibrosis gene set, indicated that the mouse model injected with CCl4 exhibited the highest similarity to human liver from NASH patients.

**Figure 6:**
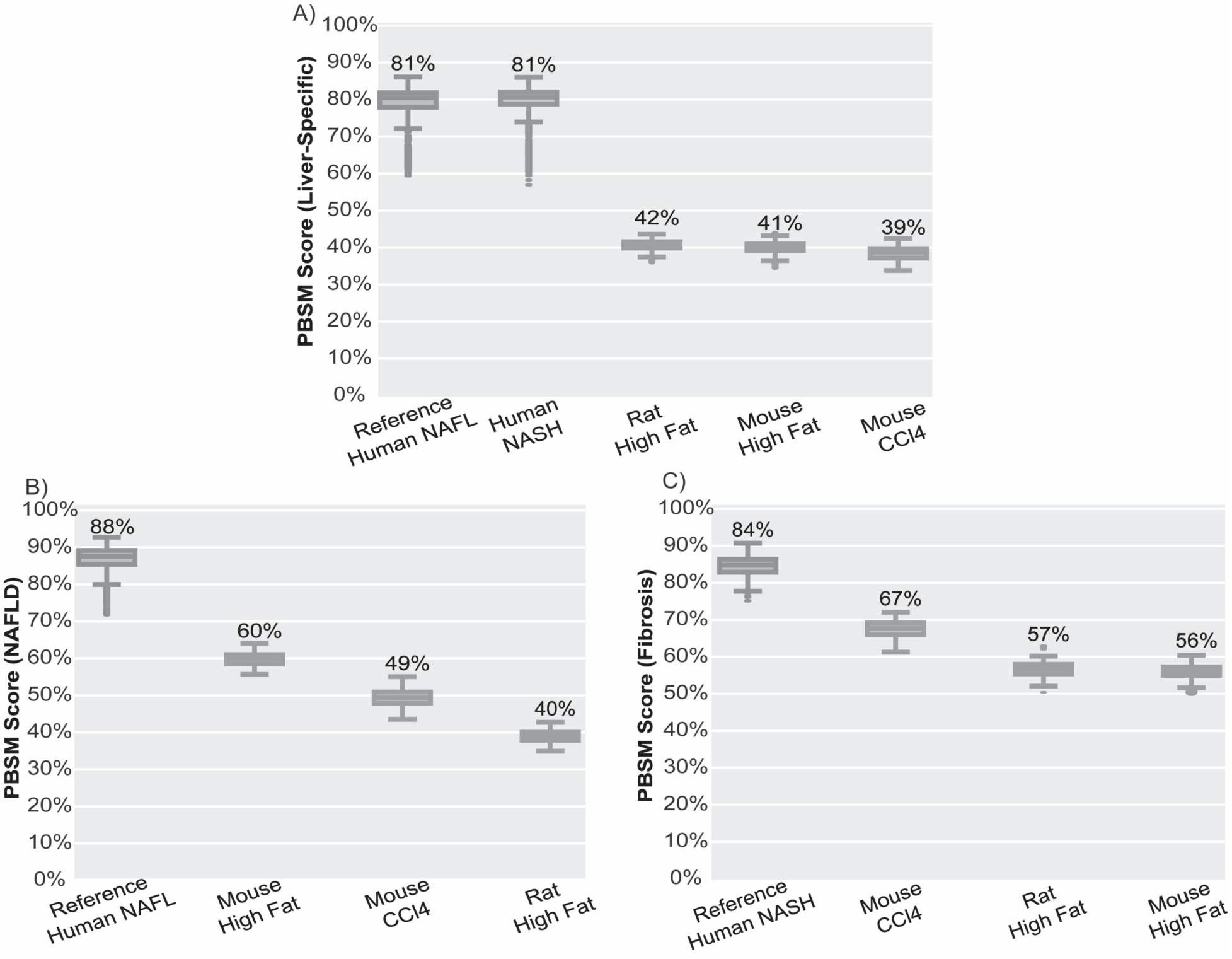
Benchmarked the transcriptomics data sets from the liver samples of various preclinical diseased animal models used in NAFLD and NASH research. A) Benchmarked human NAFL liver tissue samples using the liver-specific genes to the various diseased pre-clinical models B) Compared the pre-clinical models to the human NAFL liver tissue samples using the KEGG NAFL pathway to obtain the PBSM-based similarity scores. C) Compared the pre-clinical models to the human NASH liver tissue samples using custom fibrosis gene sets to obtain PBSM-based similarity scores.

These data underscore the critical importance of precisely defining the pathways and gene sets related to disease hallmarks when employing PBSM-based similarities for comparative analysis in disease contexts.

## Discussion

Improving preclinical prediction for drug discovery and development requires a fundamental shift toward more sophisticated and physiologically relevant human *in vitro* models. While many conventional preclinical models are widely used for various CoU applications, they also often fall short in replicating the specificity and complexity of human physiology and lead to unreliable predictions of drug efficacy and safety in humans. Additionally, the emergence of drug targets and therapeutic modalities characterized by increased human specificity, such as mRNA-based therapeutics, further accentuates the need for testing new drug candidates on more human-relevant models to mitigate the translational gap. Physiologically relevant human *in vitro* models, such as 2D and 3D primary and iPSC cultures, organoids, and MPS, offer a promising solution as alternative models for many preclinical CoU studies. However, identifying predictive CoUs for both existing and novel preclinical models requires testing a large number of test compounds for an intended CoU, which is an empirical and resource-intensive methodology with a low success rate. To expedite the identification of human physiologically relevant models for preclinical drug testing, we developed the PBSM workflow. PBSM offers a quantitative means to compare *in vitro* and *in vivo* systems, enabling researchers to assess the relative similarity between these model systems and human organs or tissues. This quantitative approach provides a numerical basis for evaluating the physiological relevance of *in vitro* and *in vivo* models. Researchers can customize PBSM analyses by selecting specific pathways or gene sets relevant to their research questions. This customization allows for a targeted evaluation of key biological processes, increasing the specificity of the assessment.

*In vitro* systems inherently simplify the complexity of *in vivo* environments. While PBSM can quantify similarities, it cannot fully overcome the limitations imposed by this simplification. Therefore, the results may not capture all aspects of the *in vivo* physiology. By incorporating data from multiple donors, PBSM can capture a broader spectrum of genetic and physiological variations, increasing the robustness and generalizability of the analysis. The sample size exerts a considerable influence on the outcomes derived from PBSM workflows, impacting the reliability of the ensuing statistical analyses. Larger sample sizes tend to yield more statistically robust results, which may pose challenges, particularly when working with limited tissue samples. Moreover, given its focus on gene-level analysis, PBSM may not be the optimal choice for analyzing alternative splicing or conducting analyses at the isoform level. The size of the pathways and gene sets under examination assumes significance in our analysis; nevertheless, the number of genes for a specific pathway introduces a critical dimension that necessitates careful consideration since the number of genes is directly correlated with the statistical confidence of the results. This interplay between pathway size and the PBSM results warrants further exploration in future investigations.

In preclinical ADME studies, various *in vitro* cultures have been used to estimate pharmacokinetic parameters for new chemical entities. The industry standard for hepatic clearance studies is the incubation of test compounds for short period of time (∼4 h) with human primary hepatocytes in suspension before cells lose it functionality due to lack of cellular attachment. The plated culture of primary human hepatocytes, iPSC-derived hepatocytes, and HepG2 cells allows longer culture periods, but these cultures are less metabolically active than suspension hepatocytes[34][35][36] and hence less suitable for *in vitro* drug metabolism studies. The PBSM results using an ADME-specific gene set demonstrated similar trends among four studies culture types. Suspension PHH has the highest PBSM value, which was comparable to human liver reference, whereas other hepatic cultures had low PBSM values. Additionally, PBSM analysis can be used to compare the effect of various *in vitro* culture methods, such as suspension versus plated *in vitro* cultures or 2D vs 3D cultures.

Cross-species differences between preclinical animal models and humans are one of the main challenges to extrapolate animal PK to human PK data[37][38][39]. The factor contributing the cross-species differences in PK predictions include differences in drug metabolizing enzymes (both abundance and activity), transporters, ADME pathways, and other physiological differences such as organ sizes, blood flow rates, and bile composition[28][40][41]. While these factors have confounding effects on species differences in ADME and cannot be studied in isolation, in this study we used PBSM analysis with ADME gene sets (including drug metabolizing enzymes and transporters) to quantitate the liver-specific cross species differences based on gene expression signatures. The liver samples from the mouse model resulted in the highest similarity score to human liver samples, while samples from the rat model had the lowest score. However, more comprehensive evaluation of animal models using PBSM for ADME applications requires inclusion of additional organs and tissues, such as gastrointestinal (GI) tract for absorption and kidney for excretion processes.

In this study, we used a modular PBSM-based similarity score methodology to study similarities of *in vitro* iPSC-CM models to human cardiac tissues using cardiac-specific gene set and a gene set for pathways involved in QT prolongation. Monitoring the maturation state and duration of iPSC-derived *in vitro* models is an essential step for quality control. Here, the PBSM analysis was applied to evaluate the effects of maturation duration and culture type (2D and 3D) of iPSC-CM models. The results demonstrate that the iPSC-CM models mature and become more physiologically similar over time. Additionally, similarity scores are sustained in 3D long-term culture of iPSC-CMs but decline over time in 2D cultures. The literature, which we obtained the iPSC-CM transcriptomics data sets from, showed phenotypical characterization of the models, suggesting that 3D iPSC-CM culture displayed improved maturation over standard 2D cultures[26][27].

Evaluation of QT prolongation using human iPSC-CM models provides predictive data to assess drug-induced pro-arrhythmia risk in humans[28][42][43]. The PBSM-based similarity scores for gene sets associated in QT prolongation was calculated for iPSC-CM models from two different iPSC-cell lines with two different culture methods (2D and 3D). While the positive predictive value (PPV) of each model still needs to be assessed empirically using test compounds, this analysis revealed the quantitative differences of various *in vitro* models that can be used for QT prolongation assay. The Cor.4u model that resulted in the highest similarity score has been assessed rigorously by the FDA for its PPV and reproducibility. Thus, use of iPSC-CMs for QT prolongation predictions under a comprehensive *in vitro* proarrhythmia assay (CiPA) is feasible[28][42][43]. We believe that the PBSM method could be used to assess the physiological relevance of *in vitro* models for narrowing down potential models that will be further qualified with CoU-specific experimental testing with large number of test compounds.

Although the pharmacodynamics of drug candidates are verified with animal disease models, over 50% of the clinical attritions during clinical phase 2 and 3 trials are due to lack of efficacy[44]. Recapitulation of human disease in animals, especially chronic diseases with various genetic risk factors, such as metabolic dysfunction-associated fatty liver disease and NASH, poses several challenges due to the differences in etiology, complexity, and progression of pathophysiologies between animal models and humans. In this study, PBSM-based similarity scores were calculated for three conventional animal models used to study NAFLD and NASH disease stages [45][ [46][47][48] to compare human liver-specific, NAFLD, and NASH-specific pathways. Overall, the similarity scores of each animal model were significantly lower than the variability observed in human reference samples. This suggests that animal models offer poor representations of human diseases with respect to organ-specific gene expression profiles. However, PBSM analysis further revealed higher similarity scores for the commonly used animal models, such as mouse model with high-fat diet for NAFLD modeling [49] and mouse model with CCl4 induced liver fibrosis modeling [50] compared to other models including rat disease models.

In this work, we demonstrated that PBSM can be applied in the early stages of research to evaluate the suitability of *in vitro* and *in vivo* models for specific CoU applications. This can help researchers identify promising models for further investigation and prioritize resources efficiently. PBSM provides an objective and data-driven assessment of *in vitro* and *in vivo* systems, reducing the reliance on subjective judgments when selecting or validating models for particular CoUs. These analyses support that PBSM can be applied at various stages of NAM development and qualification studies along with phenotypic data (e.g., metabolomic and proteomic profiling) for more comprehensive and functional comparison of preclinical models and humans.

## Study Highlights

### What is the current knowledge on the topic?

Translational Gap: While preclinical studies frequently yield encouraging results in cellular and animal models, these findings may not always translate into clinical efficacy and safety in human subjects due to species-specific differences and heterogeneity among individual patients. Current methodologies for evaluating the human relevance of preclinical models rely on comparing a limited number of qualitative and quantitative metrics, such as morphology, phenotypical metrics, and responses to known test compounds, and hence those do not provide a comprehensive evaluation for human-relevancy.

### What question did this study address?

This study introduces the Pathway-Based Similarity Measurement (PBSM) as a novel approach to quantify transcriptomic similarity between distinct biological samples. PBSM assesses the similarity between biological samples by performing pairwise comparison (e.g. human vs animal tissues) based on user-defined gene sets. PBSM holds the potential to evaluate human relevance of preclinical models and new approach method (NAMs) for intended context of use (CoU) application.

### What does this study add to our knowledge?

This study offers a quantitative method to compare the transcriptomic similarity of preclinical models to human tissues, shown here as proof of concept for liver and cardiac tissues, enabling improved model selection and validation. PBSM can successfully support CoU selection for preclinical models, assess the impact of different gene sets on similarity calculations, and differentiate among various in vitro and in vivo models.

### How might this change drug discovery, development, and/or therapeutics?

This study presents a novel and versatile approach for addressing the unmet need for quantitative assessment of human relevance of preclinical models. This approach can be used in conjunction with the current methodologies for qualification of preclinical models and may accelerate the development of human-relevant models, which is essential to bridge the translational gap between preclinical and clinical studies.

## Acknowledgements

The authors thank Dr. Brian Berridge for providing invaluable guidance and discussions during the manuscript review.

The authors would like to thank the Genotype-Tissue Expression (GTEx) Project which was supported by the Common Fund of the Office of the Director of the National Institutes of Health, and by NCI, NHGRI, NHLBI, NIDA, NIMH, and NINDS. The data used for the analyses described in this manuscript were obtained from the GTEx Portal on 12/22/2022.

## Author Contributions

P.P., M.C., and J.S. wrote the manuscript; M.C., B.A., and P.P. designed the research; P.P. and J.S. performed the research; P.P. analyzed the data.

## Code Availability

The code for PBSM has been implemented in R and can be downloaded with example datasets from https://github.com/JavelinBiotech/Pathway-Based-similarity/.

## References

1. Conesa A., Madrigal P., Tarazona S., et al. A survey of best practices for RNA-seq data analysis. Genome Biol 2016; 17; 13. doi: 10.1186/s13059-016-0881-8

2. Love M.I., Huber W. & Anders S. Moderated estimation of fold change and dispersion for RNA-seq data with DESeq2. Genome Biol 2014; 15; 550. doi: 10.1186/s13059-014-0550-8

3. Radley A., Schwab R., Tan Y., et al. Assessment of engineered cells using CellNet and RNA-seq. Nat Protoc 2017;12;1089–1102. doi: 10.1038/nprot.2017.022

4. Li W.V., Chen Y. & Li J.J. TROM: A Testing-Based Method for Finding Transcriptomic Similarity of Biological Samples. Stat Biosci 2017;9;105–136. doi: 10.1007/s12561-016-9163-y

5. Kim D., Ryu J., Son M., et al. A liver-specific gene expression panel predicts the differentiation status of in vitro hepatocyte models. Hepatology 2017;66(5);1662–1674. doi: 10.1002/hep.29324

6. Zhao Y., Li M., Konaté M., et al. TPM, FPKM, or Normalized Counts? A Comparative Study of Quantification Measures for the Analysis of RNA-seq Data from the NCI Patient-Derived Models Repository. J Transl Med 2021;19;269. doi: 10.1186/s12967-021-02936-w

7. Kanehisa M., Goto S., Hattori M., et al. KEGG: Kyoto Encyclopedia of Genes and Genomes, Nucleic Acids Research 2000; 28; 27–30. doi: 10.1093/nar/28.1.27

8. Gillespie M., Jassal B., Stephan R., et al. The reactome pathway knowledgebase 2022, Nucleic Acids Research 2022; 50; D687–D692, doi: 10.1093/nar/gkab1028

9. Subramanian A., Tamayo P., Mootha VK., et al. Gene set enrichment analysis: a knowledge-based approach for interpreting genome-wide expression profiles. Proc Natl Acad Sci U S A. 2005;102(43);15545–50. doi: 10.1073/pnas.0506580102.

10. Mehri A., Jamaati M. & Hassan M. Word ranking in a single document by Jensen–Shannon divergence, Physics Letters A. North-Holland 2015; 379; 1627–1632. doi: 10.1016/j.physleta.2015.04.030.

11. Capra J.A., & Singh M. Predicting functionally important residues from sequence conservation. Bioinformatics. 2007;23(15):1875–82. doi: 10.1093/bioinformatics/btm270.

12. Lin J., Divergence measures based on the Shannon entropy. IEEE Transactions 1991; 37; 145–151. doi: 10.1109/18.61115

13. Sims G., Jun S., Wu G., et al. Alignment-free genome comparison with feature frequency profiles (FFP) and optimal resolutions. Proc Natl Acad Sci U S A. 2009;106(8);2677–82. doi: 10.1073/pnas.0813249106.

14. Itzkovitz S., Hodis E. & Segal E. Overlapping codes within protein-coding sequences. Genome Res. 2010; 20(11); 1582–1589. doi: 10.1101/gr.105072.110.

15. Ré M., & Azad R. Generalization of entropy based divergence measures for symbolic sequence analysis. PloS 2014; one vol. 9,4. doi: 10.1371/journal.pone.0093532

16. Hou J., Xiufen Y., Weixing F., et al. Distance correlation application to gene co-expression network analysis. BMC bioinformatics 2022; 23; doi: 10.1186/s12859-022-04609-x

17. Uhlén M., Fagerberg L., Björn M., et al. Tissue-Based Map of the Human Proteome. Science 347 2015; 6220; 1260419. doi: 10.1126/science.1260419. https://www.proteinatlas.org/.

18. Doogue M., & Polasek T. The ABCD of clinical pharmacokinetics. Therapeutic advances in drug safety vol. 2013; 4,1; 5–7. doi: 10.1177/2042098612469335

19. Han S., Park J., Lee J., et al. Targeted Next-Generation Sequencing for Comprehensive Genetic Profiling of Pharmacogenes. Clinical pharmacology and therapeutics 2017; 101; 396–405. doi: 10.1002/cpt.532

20. Boon R., Kumar M., Tricot T., et al. Amino acid levels determine metabolism and CYP450 function of hepatocytes and hepatoma cell lines. Nature communications 2020; 11; 1 1393. doi: 10.1038/s41467-020-15058-6

21. Moreau F., Brunao B., Liu X., et al. Liver-specific FGFR4 knockdown in mice on an HFD increases bile acid synthesis and improves hepatic steatosis. Journal of lipid research 2023; 64; 100324. doi: 10.1016/j.jlr.2022.100324

22. Gong B., Wang C., Su Z., et al. Transcriptomic profiling of rat liver samples in a comprehensive study design by RNA-Seq. Scientific data 2014; 1. doi: 10.1038/sdata.2014.21

23. Jin L., Tang Q., Hu S., et al. A pig BodyMap transcriptome reveals diverse tissue physiologies and evolutionary dynamics of transcription. Nature communications 2021; 12(1); 3715. doi: 10.1038/s41467-021-23560-8.

24. Cox C., Lynch S., Goldring C., et al. Current Perspective: 3D Spheroid Models Utilizing Human-Based Cells for Investigating Metabolism-Dependent Drug-Induced Liver Injury. Frontiers in medical technology 2020; 2; 611913. doi: 10.3389/fmedt.2020.611913.

25. Stransky S., Cutler R., Aguilan J., et al. Investigation of reversible histone acetylation and dynamics in gene expression regulation using 3D liver spheroid model. Epigenetics & chromatin 2022; 15(1). doi: 10.1186/s13072-022-00470-7.

26. Branco M., Cotovio J., Rodrigues C., et al. Transcriptomic analysis of 3D Cardiac Differentiation of Human Induced Pluripotent Stem Cells Reveals Faster Cardiomyocyte Maturation Compared to 2D Culture. Scientific reports 2019; 9(1); 9229. doi: 10.1038/s41598-019-45047-9

27. Ergir E., Cruz J., Fernandes S., et al. Generation and maturation of human iPSC-derived 3D organotypic cardiac microtissues in long-term culture. Scientific reports 2022; 12(1); 17409. doi: 10.1038/s41598-022-22225-w

28. Blinova K., Stohlman J., Vicente J., et al. Comprehensive Translational Assessment of Human-Induced Pluripotent Stem Cell Derived Cardiomyocytes for Evaluating Drug-Induced Arrhythmias. Toxicological sciences: an official journal of the Society of Toxicology 2017; vol. 155(1); 234–247. doi:10.1093/toxsci/kfw200

29. Wang H., Sheehan R., Palmer A., et al. Adaptation of Human iPSC-Derived Cardiomyocytes to Tyrosine Kinase Inhibitors Reduces Acute Cardiotoxicity via Metabolic Reprogramming. Cell systems 2019; 8(5); 412–426. doi: 10.1016/j.cels.2019.03.009

30. Loewa A., Feng J., & Hedtrich S. Human disease models in drug development. Nature reviews bioengineering 2023; 1–15. doi: 10.1038/s44222-023-00063-3

31. Kozumi K., Kodama T., Murai H., et al. Transcriptomics Identify Thrombospondin-2 as a Biomarker for NASH and Advanced Liver Fibrosis. Hepatology (Baltimore, Md.) 2021; 74(5); 2452–2466. doi: 10.1002/hep.31995.

32. Xi Y., LaCanna R, Ma H.Y., et al. A WISP1 antibody inhibits MRTF signaling to prevent the progression of established liver fibrosis. Cell metabolism 2022; 34(9); 1377–1393. doi: 10.1016/j.cmet.2022.07.009.

33. Metzner V., Herzog G., Heckel T., et al. Liraglutide + PYY3-36 Combination Therapy Mimics Effects of Roux-en-Y Bypass on Early NAFLD Whilst Lacking-Behind in Metabolic Improvements. Journal of clinical medicine 2022; 11(3); 753. doi: 10.3390/jcm11030753.

34. Kratochwil NA, Meille C, Fowler S, et al. Metabolic Profiling of Human Long-Term Liver Models and Hepatic Clearance Predictions from In Vitro Data Using Nonlinear Mixed-Effects Modeling. AAPS J. 2017; 19(2); 534–550. doi: 10.1208/s12248-016-0019-7.

35. Gerets HH, Tilmant K, Gerin B, et al. Characterization of primary human hepatocytes, HepG2 cells, and HepaRG cells at the mRNA level and CYP activity in response to inducers and their predictivity for the detection of human hepatotoxins. Cell Biol Toxicol. 2012; 28(2); 69–87. doi: 10.1007/s10565-011-9208-4.

36. Kvist A.J., Kanebratt K.P., Walentinsson A., et al. Critical differences in drug metabolic properties of human hepatic cellular models, including primary human hepatocytes, stem cell derived hepatocytes, and hepatoma cell lines. Biochem Pharmacol. 2018; 155; 124–140. doi: 10.1016/j.bcp.2018.06.026.

37. Chu X., Bleasby K. & Evers R. Species differences in drug transporters and implications for translating preclinical findings to humans. Expert Opin Drug Metab Toxicol. 2013; 9(3); 237–52. doi: 10.1517/17425255.2013.741589.

38. Baillie T.A., & Rettie A.E. Role of biotransformation in drug-induced toxicity: influence of intra- and inter-species differences in drug metabolism. Drug Metab Pharmacokinet. 2011; 26(1); 15–29. doi: 10.2133/dmpk.dmpk-10-rv-089.

39. Dalgaard L. Comparison of minipig, dog, monkey and human drug metabolism and disposition. J Pharmacol Toxicol Methods. 2015; 74; 80–92. doi: 10.1016/j.vascn.2014.12.005.

40. Martinez M.N., Mochel J.P., Neuhoff S., et al. Comparison of Canine and Human Physiological Factors: Understanding Interspecies Differences that Impact Drug Pharmacokinetics. AAPS J. 2021; 23(3); 59. doi: 10.1208/s12248-021-00590-0.

41. Martignoni M., Groothuis G. & Kanter R. Species differences between mouse, rat, dog, monkey and human CYP-mediated drug metabolism, inhibition and induction. Expert Opin Drug Metab Toxicol. 2006; 2(6); 875–94. doi: 10.1517/17425255.2.6.875.

42. Lu HR., Zeng H., Kettenhofen R., et al. Assessing Drug-Induced Long QT and Proarrhythmic Risk Using Human Stem-Cell-Derived Cardiomyocytes in a Ca2+ Imaging Assay: Evaluation of 28 CiPA Compounds at Three Test Sites. Toxicol Sci. 2019; 170(2); 345–356. doi: 10.1093/toxsci/kfz102.

43. Blinova K., Dang Q., Millard D., et al. International Multisite Study of Human-Induced Pluripotent Stem Cell-Derived Cardiomyocytes for Drug Proarrhythmic Potential Assessment. Cell Rep. 2018; 24(13); 3582–3592. doi: 10.1016/j.celrep.2018.08.079.

44. Arrowsmith J., & Miller P. Trial watch: phase II and phase III attrition rates 2011-2012. Nat Rev Drug Discov. 2013; 12(8); 569. doi: 10.1038/nrd4090.

45. Ipsen D.H., Lykkesfeldt J. & Tveden-Nyborg P.. Animal Models of Fibrosis in Nonalcoholic Steatohepatitis: Do They Reflect Human Disease? Adv Nutr. 2020; 11(6); 1696–1711. doi: 10.1093/advances/nmaa081.

46. Peng C., Stewart A., Woodman O.L., et al. Non-Alcoholic Steatohepatitis: A Review of Its Mechanism, Models and Medical Treatments. Front Pharmacol. 2020; 11; 603926. doi: 10.3389/fphar.2020.603926.

47. Hansen H., Feigh M., Veidal S.S., et al. Mouse models of nonalcoholic steatohepatitis in preclinical drug development. Drug Discov Today. 2017; 22(11); 1707–1718. doi: 10.1016/j.drudis.2017.06.007.

48. Bao Y.L., Wang L., Pan H., et al. Animal and Organoid Models of Liver Fibrosis. Front Physiol. 2021; 12; 666138. doi: 10.3389/fphys.2021.666138.

49. Chen X., Acquaah-Mensah G., Denning K.L., et al. High-fat diet induces fibrosis in mice lacking CYP2A5 and PPARα: a new model for steatohepatitis-associated fibrosis. Am J Physiol Gastrointest Liver Physiol. 2020; 319(5); G626–G635. doi: 10.1152/ajpgi.00213.2020.

50. Scholten D., Trebicka J., Liedtke C., et al. The carbon tetrachloride model in mice. Lab Anim. 2015; 49; 4–11. doi: 10.1177/0023677215571192.

