## Supplemental Files for "Pathway-based similarity measurement to quantify transcriptomics similarity between human tissues and preclinical models"

Supplementary Information for the manuscript **Pathway-based similarity measurement to quantify transcriptomics similarity between human tissues and preclinical models.**

### Supplementary Materials & Methods

#### <u>Data Sources and Processing</u>

The RNA-seq datasets for human tissue samples were obtained from the Genotype-Tissue Expression (GTEx) database by the Broad Institute. Samples were initially filtered for the donors with liver tissue samples. Additionally, the sub-samples were omitted from the analysis (Esophagus, Cervix, Minor Salivary Gland, Fallopian Tube, etc). Based on this criteria, a total of 226 donors, 4163 samples were used (Figure S1).

A subset of 226 donors with a total of 1339 samples including biopsies of the Heart, Brain, Colon, Kidney, Liver, Muscle, and Small Intestine were used for further analysis.

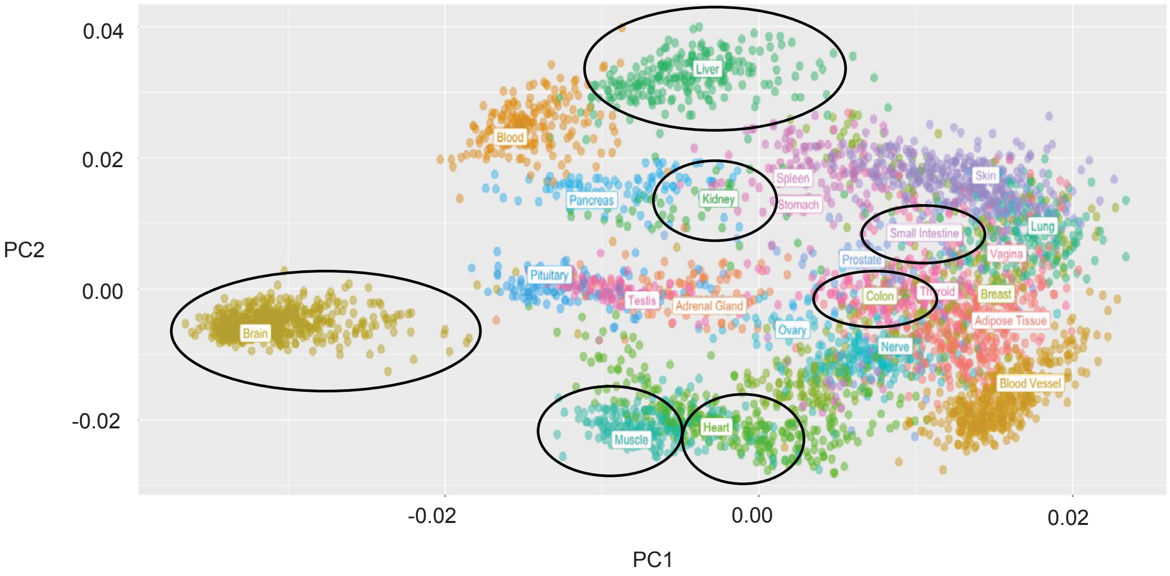

*Figure S1:* GTEX data samples distribution using PCA after processing.

The RNA-seq transcriptomics datasets from GEO used in these studies pre-processed to generate the gene counts. Pre-processing the data involved quality checks using fastqc [1], trimming of the data to remove adapters using fastp [2], and the gene counts were then generated using salmon [3]. For the animal RNA-seq transcriptomics data, each dataset went through the same pipeline, but orthologs were generated for the animal genes to human genes using biomart [4]. Further processing was conducted to obtain the transcripts per million (TPMs) from the raw reads.

**PBSM Input**

PBSM uses the normalized gene counts and the gene sets for comparison. The normalized gene counts are obtained from the pre-processing pipeline as described earlier, and the gene sets are defined from the pathway databases as well as custom gene sets obtained from different literature sources. These gene sets for ADME analysis have been defined in Table S1, Table S2, and Table S3. Further, the gene sets for the Cardiac Maturation, QT prolongation, and Fibrosis have been defined in Table S4, Table S5, and Table S6 respectively.

*Table S3: Summary of the gene set used for the PBSM calculations for the genes involved in ADME as defined in* Han et al., 2016 [8].

| | | | | | | | | |
|---|---|---|---|---|---|---|---|---|
| ABCA1 | AHR | CYP1A2 | CYP7A1 | GRK5 | NAT2 | PTGS2 | SLC47A1 | UGT1A1 |
| ABCB1 | ALDH1A1 | CYP2A6 | DBH | GSTM1 | NQO1 | RYR1 | SLC47A2 | UGT1A4 |
| ABCB11 | ALOX5 | CYP2B6 | DPYD | GSTP1 | NR1I2 | RYR2 | SLC6A3 | UGT1A9 |
| ABCC2 | APOA1 | CYP2C19 | DRD1 | GSTT1 | NR1I3 | SCN1A | SLC6A4 | UGT2B15 |
| ABCC4 | ARID5B | CYP2C8 | DRD2 | HMGCR | NR3C2 | SCN2A | SLCO1A2 | UGT2B7 |
| ABCG1 | BDNF | CYP2C9 | EGFR | HTR1A | NTRK2 | SCN5A | SLCO1B1 | VDR |
| ABCG2 | BRCA1 | CYP2D6 | EPHX1 | HTR2A | P2RY1 | SLC15A1 | SLCO1B3 | VKORC1 |
| ACE | CES1 | CYP2E1 | ESR1 | KCNH2 | P2RY12 | SLC15A2 | SLCO2B1 | ZNF423 |
| ADH1A | CES2 | CYP2J2 | F5 | KCNJ11 | PEAR1 | SLC19A1 | SOD2 | |
| ADH1B | COMT | CYP2R1 | FKBP5 | LDLR | PON1 | SLC22A1 | SULT1A1 | |
| ADH1C | CPS1 | CYP3A4 | G6PD | MAOA | POR | SLC22A2 | TBXAS1 | |
| ADRB1 | CRHR1 | CYP3A5 | GLCCI1 | MTHFR | PTGIS | SLC22A3 | TPMT | |
| ADRB2 | CYP1A1 | CYP4F2 | GRK4 | NAT1 | PTGS1 | SLC22A6 | TYMS | |

Table S1 consists of a list of genes involved in ADME metabolism as defined in Han et al., 2016 [8]. Pairwise PBSM calculations were performed using this dataset in Figure 2a, Figure 4a, Figure 4b, and Figure 4c.

*Table S2: Summary of the gene set used for the PBSM calculations for the genes involved in ADME as defined by the KEGG Database. (5)(6)*

| | | | | | | | |
|---|---|---|---|---|---|---|---|
| ADH1A | CES2 | CYP3A4 | GSTA2 | GSTT1 | NAT1 | UGT1A5 | UGT2B7 |
| ADH1B | CES5A | CYP3A43 | GSTA3 | GSTT2 | NAT2 | UGT1A6 | UMPS |
| ADH1C | CYP1A2 | CYP3A5 | GSTA4 | GSTZ1 | TK1 | UGT1A7 | UPB1 |
| ADH4 | CYP2A13 | CYP3A7 | GSTA5 | GUSB | TK2 | UGT1A8 | UPP1 |
| ADH5 | CYP2A6 | DPYD | GSTK1 | HPRT1 | TPMT | UGT1A9 | UPP2 |
| ADH6 | CYP2A7 | DPYS | GSTM1 | IMPDH1 | TYMP | UGT2A1 | XDH |
| ADH7 | CYP2B6 | FMO1 | GSTM2 | IMPDH2 | UCK1 | UGT2A3 | |
| ALDH1A3 | CYP2C18 | FMO2 | GSTM3 | ITPA | UCK2 | UGT2B10 | |
| ALDH3A1 | CYP2C19 | FMO3 | GSTM4 | MAOA | UCKL1 | UGT2B11 | |
| ALDH3B1 | CYP2C8 | FMO4 | GSTM5 | MAOB | UGT1A1 | UGT2B15 | |
| ALDH3B2 | CYP2C9 | FMO5 | GSTO1 | MGST1 | UGT1A10 | UGT2B17 | |
| AOX1 | CYP2D6 | GMPS | GSTO2 | MGST2 | UGT1A3 | UGT2B28 | |
| CDA | CYP2E1 | GSTA1 | GSTP1 | MGST3 | UGT1A4 | UGT2B4 | |

Table S2 consists of a list of genes involved in ADME metabolism as defined by the KEGG database. This gene set was used in the pairwise PBSM calculations to obtain the median similarity scores for the different GTEX tissue samples defined in Figure 2a.

*Table S3: Summary of the gene set used for the PBSM calculations for the genes involved in ADME as defined by Reactome Database. (7)*

| | | | | | | | |
|---|---|---|---|---|---|---|---|
| ABCB1 | ADH1A | GGT1 | GSTT1 | PON3 | SLCO2B1 | UGT1A6 | UGT2B7 |
| ABCC1 | ALB | GGT5 | GUK1 | RAC1 | SULT1A1 | UGT1A7 | UGT3A1 |
| ABCC2 | BCHE | GGT6 | HPRT1 | SLC16A1 | SULT1A3 | UGT1A8 | UGT3A2 |
| ABCC3 | BSG | GGT7 | IMPDH1 | SLC22A1 | SULT1A4 | UGT1A9 | VAV1 |
| ABCC4 | CES1 | GLYAT | IMPDH2 | SLC22A2 | SULT1C4 | UGT2A1 | VAV2 |
| ABCC5 | CES2 | GLYATL1 | NAT1 | SLC22A3 | SULT1E1 | UGT2A2 | VAV3 |
| ABCG2 | CNDP2 | GLYATL2 | NAT2 | SLC22A7 | SULT2A1 | UGT2A3 | XDH |

| | | | | | | |
|---|---|---|---|---|---|---|
| ACSM2A | CYP2C19 | GLYATL3 | NME1 | SLC28A2 | TPMT | UGT2B10 |
| ACSM2B | CYP2C8 | GMPS | NME2 | SLC28A3 | UGT1A1 | UGT2B11 |
| ACSM4 | CYP2C9 | GSTA1 | NT5C2 | SLC29A1 | UGT1A10 | UGT2B15 |
| ACSM5 | CYP2D6 | GSTA2 | NUDT15 | SLC29A2 | UGT1A3 | UGT2B17 |
| ACY1 | CYP2E1 | GSTM1 | PCK1 | SLCO1B1 | UGT1A4 | UGT2B28 |
| ADAL | CYP3A4 | GSTP1 | PON1 | SLCO1B3 | UGT1A5 | UGT2B4 |

Table S3 consists of a list of genes involved in ADME metabolism as defined by the Reactome database. This gene set was used in the pairwise PBSM calculations to obtain the median similarity scores for the different GTEX tissue samples defined in Figure 2a.

*Table S4: Summary of the custom gene set used for the PBSM calculations for the genes involved in cardiac maturation as defined in the Paper [11].*

| | | | | | | | | |
|---|---|---|---|---|---|---|---|---|
| ACAA1 | CACNA1I | DKK1 | GJA1 | JUN | MYH11 | PITX2 | SLC27A6 | TNNT2 |
| ACADM | CAMK2B | DKK3 | GJA5 | KCNA4 | MYH6 | PKM | SLC2A1 | TP53 |
| ACAT1 | CASQ2 | DVL1 | GJD3 | KCNA5 | MYH7 | PLN | SLC2A3 | TPI1 |
| ACO2 | CAV3 | ENO3 | GPC2 | KCND2 | MYH7B | PPARA | SLC2A4 | TRDN |
| ACSL1 | CCND1 | EOMES | GPI | KCND3 | MYL2 | PPARG | SLC8A1 | VCAM1 |
| ACTA2 | CCND2 | ETV2 | GSC | KCNH2 | MYL3 | PPARGC1A | SMAD1 | VEGFA |
| ACTC1 | CCNL1 | FABP3 | GSK3B | KCNJ12 | MYL4 | PPP3CC | SMAD2 | VEGFB |
| ACTN2 | CD34 | FGF10 | HADHA | KCNJ2 | MYL7 | PRICKLE1 | SMAD3 | VEGFC |
| ADAM10 | CD36 | FGF2 | HADHB | KCNJ3 | MYLK3 | PRKACA | SMAD5 | VIM |
| ADRB1 | CDH1 | FGF4 | HAND2 | KCNJ4 | MYOM2 | PRKD1 | SMAD6 | WNT10B |
| ADRB2 | CDH2 | FGF8 | HCN1 | KCNJ5 | MYOZ2 | PYGO1 | SMAD7 | WNT11 |
| AKAP6 | CDH5 | FGFR2 | HCN2 | KCNQ1 | NKD1 | RARB | SNAI1 | WNT2 |
| ALDOA | CDKN2A | FGFR3 | HCN4 | KDR | NKX2-5 | RB1 | SNAI2 | WNT2B |
| ANGPT1 | CDX2 | FH | HES1 | LDHA | NODAL | RBM20 | SOX17 | WNT3A |
| APC | CEBPA | FOXC1 | HEY1 | LEF1 | NOG | ROR2 | SUCLG1 | WNT4 |
| ATP2A2 | CEBPB | FOXF1 | HK1 | LIN28B | NOTCH1 | RRAS | TALDO1 | WNT5A |
| AXIN1 | CKMT2 | FRZB | HK2 | MAPK1 | NOTCH3 | RUNX1T1 | TBX5 | WNT5B |
| AXIN2 | CNN1 | FXYD1 | HMGA1 | MDH2 | NPPA | RXRA | TCAP | WNT8A |
| BCL9 | COL1A1 | FZD1 | HOXB2 | ME1 | NPPB | RYR2 | TCF4 | ZBTB16 |
| BIN1 | COL3A1 | FZD2 | HSP90AA1 | MEF2C | NRP1 | SCN1A | TCF7 | ZEB2 |
| BMP1 | CPT1B | FZD4 | IDH3A | MEIS1 | OGDH | SCN2A | TCF7L2 | ZFPM1 |
| BMP2 | CS | FZD6 | IGF2BP2 | MEIS2 | PBX3 | SCN4A | TFAM | ZNF503 |
| BMP4 | CTNNB1 | FZD7 | INHBA | MESP1 | PDGFRA | SCN5A | TGFB2 | |
| BMP5 | CXXC4 | FZD8 | IRX3 | MESP2 | PDHA1 | SDHA | TKT | |
| BMP7 | CYP2J2 | G6PD | IRX5 | MIXL1 | PECAM1 | SFRP1 | TMEM88 | |
| CACNA1C | DACT1 | GATA4 | ISL1 | MPC1 | PFKM | SFRP2 | TNNC1 | |
| CACNA1D | DACT3 | GATA5 | ITGA3 | MPC2 | PGAM2 | SFRP5 | TNNI1 | |
| CACNA1G | DCN | GATA6 | ITPR3 | MSX1 | PGK1 | SLC25A20 | TNNI3 | |
| CACNA1H | DDR2 | GDF15 | JPH2 | MYC | PGM1 | SLC27A1 | TNNI3K | |

Table S4 consists of a list of genes involved in cardiac maturation as defined in the Paper [31239450]. This gene set was used in the pairwise PBSM calculations to obtain the median similarity scores for the maturation over time for the 2D and 3D iPSC-derived cardiomyocytes [Figure 5a].

*Table S*5: Summary of the custom gene set used for the PBSM calculations for the genes involved in QT prolongation.

| | | | | | | | | |
|---|---|---|---|---|---|---|---|---|
| ABCB1 | ATP2A2 | CASQ2 | KCNA2 | KCNH2 | KCNJ6 | KCNMB4 | RYR2 | SCN5A |
| ABCC9 | CACNA1C | CAV3 | KCNA5 | KCNH6 | KCNJ8 | KCNN3 | SCN10A | SCN9A |
| AKAP6 | CACNB1 | CERKL | KCNB1 | KCNH7 | KCNK2 | KCNQ1 | SCN11A | SNTA1 |
| AKAP7 | CACNB2 | CNOT3 | KCNB2 | KCNJ11 | KCNK3 | MINK1 | SCN1B | TRDN |
| AKAP9 | CALM1 | GPD1L | KCND3 | KCNJ12 | KCNMA1 | NOS1AP | SCN2B | ZFHX3 |
| ANK2 | CALM2 | JPH2 | KCNE1 | KCNJ2 | KCNMB1 | NUBPL | SCN3B | |
| APLP2 | CALM3 | JPH3 | KCNE2 | KCNJ3 | KCNMB2 | PLN | SCN4A | |
| ATP1B1 | CALR | KCNA1 | KCNH1 | KCNJ5 | KCNMB3 | PPP2R3A | SCN4B | |

Table S5 consists of a list of genes involved in QT prolongation. This gene set was used in the pairwise PBSM calculations to obtain the median similarity scores corresponding to the different iPSC model's ability to predict drug-induced QT-prolongation [Figure 5c]

*Table S*6: Summary of the gene set used for the PBSM calculations for the genes defined in the KEGG NAFLD Pathway.[13]

| | | | | | | | |
|---|---|---|---|---|---|---|---|
| ADIPOQ | COX5A | CYCS | INSR | MLXIP | NDUFB1 | NDUFS4 | PPARA |
| ADIPOR1 | COX5B | CYP2E1 | IRS1 | MLXIPL | NDUFB10 | NDUFS5 | PPARG |
| ADIPOR2 | COX6A1 | DDIT3 | IRS2 | NDUFA1 | NDUFB11 | NDUFS6 | PRKAA1 |
| AKT1 | COX6A2 | EIF2AK3 | ITCH | NDUFA10 | NDUFB2 | NDUFS7 | PRKAA2 |
| AKT2 | COX6B1 | EIF2S1 | JUN | NDUFA11 | NDUFB3 | NDUFS8 | PRKAB1 |
| AKT3 | COX6B2 | ERN1 | LEP | NDUFA12 | NDUFB4 | NDUFV1 | PRKAB2 |
| ATF4 | COX6C | FAS | LEPR | NDUFA13 | NDUFB5 | NDUFV2 | PRKAG1 |
| BAX | COX7A1 | FASLG | MAP3K11 | NDUFA2 | NDUFB6 | NDUFV3 | PRKAG2 |
| BCL2L11 | COX7A2 | FOS | MAP3K5 | NDUFA3 | NDUFB7 | NFKB1 | PRKAG3 |
| BID | COX7A2L | GSK3A | MAPK10 | NDUFA4 | NDUFB8 | NR1H3 | RAC1 |
| CASP3 | COX7B | GSK3B | MAPK11 | NDUFA4L2 | NDUFB9 | PIK3CA | RELA |
| CASP7 | COX7B2 | IKBKB | MAPK12 | NDUFA5 | NDUFC1 | PIK3CB | RXRA |
| CASP8 | COX7C | IL1A | MAPK13 | NDUFA6 | NDUFC2 | PIK3CD | SDHA |
| CDC42 | COX8A | IL1B | MAPK14 | NDUFA7 | KCTD14 | PIK3R1 | SDHB |
| CEBPA | COX8C | IL6 | MAPK8 | NDUFA8 | NDUFS1 | PIK3R2 | SDHC |
| COX4I1 | CXCL8 | IL6R | MAPK9 | NDUFA9 | NDUFS2 | PIK3R3 | SDHD |
| COX4I2 | CYC1 | INS | MLX | NDUFAB1 | NDUFS3 | PKLR | SOCS3 |
| SREBF1 | TNFRSF1A | TNF | XBP1 | UQCRB | UQCRC2 | UQCRH | UQCRQ |
| TGFB1 | TRAF2 | UQCR10 | UQCR11 | UQCRC1 | UQCRFS1 | UQCRHL | |

Table S6 consists of a list of genes defined by the KEGG database for the NAFLD pathway. This gene set was used in the pairwise PBSM calculations to obtain the median similarity scores for the different *in-vivo* models used to model pre-clinical NAFLD disease [Figure 6b].

*Table S*7: Summary of the custom gene set used for the PBSM calculations for the genes involved in Fibrosis.

| | | | | | | | | |
|---|---|---|---|---|---|---|---|---|
| ACAT2 | CCL4 | COL5A1 | EGR1 | HMGCR | LOXL2 | PCK1 | SMAD3 | TNF |
| APOA1 | CCL5 | COL6A1 | FASN | IGF1 | MAPK1 | PINK1 | SOD2 | XBP1 |
| APOC3 | CCR2 | COL6A2 | FGF2 | IGF1R | MAPK14 | PPARA | SPP1 | |
| APOE | CCR5 | COL6A3 | FGF7 | IL10 | MAPK3 | PPARD | STAT3 | |
| ATF4 | COL1A1 | CXCL10 | GCK | IL1B | MMP13 | PPARG | TGFB1 | |
| CAT | COL1A2 | CYP2E1 | GLI2 | IL22 | MMP2 | PTEN | TIMP1 | |
| CCL2 | COL3A1 | DGAT2 | GLI3 | IL6 | MMP9 | PTPN1 | TIMP2 | |
| CCL20 | COL4A1 | EGF | HGF | KEAP1 | NLRP3 | SHH | TIMP3 | |
| CCL3 | COL4A2 | EGFR | HIF1A | LOX | NRF1 | SIRT1 | TLR4 | |

Table S7 consists of a list of genes involved in liver Fibrosis. This gene set was used in the pairwise PBSM calculations to obtain the median similarity scores for the different *in-vivo* diseased models used to model pre-clinical liver fibrosis [Figure 6c].

### PBSM Supplementary Results

Black and White Figures:

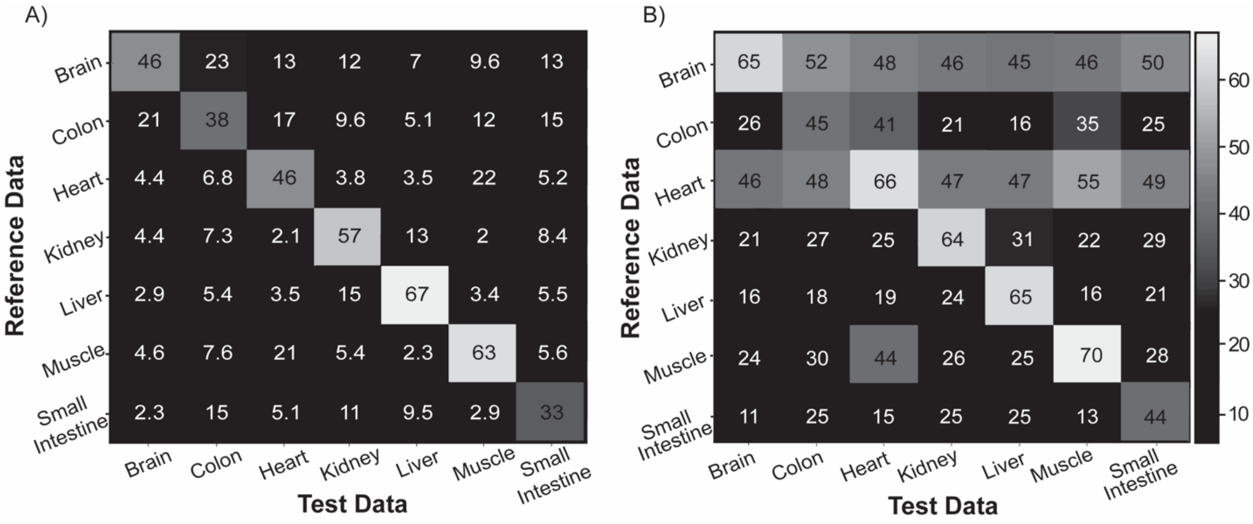

*Figure 2 (B&W):* Assessment of tissue-specificity using the PBSM algorithm for the organ-specific & highly expressed gene sets. A) Median similarity scores that are obtained using the organ-specific gene set across the different RTS and TTS tissue samples. B) Median similarity scores that were obtained for the highly expressed gene set across the different RTS and TTS tissue samples.

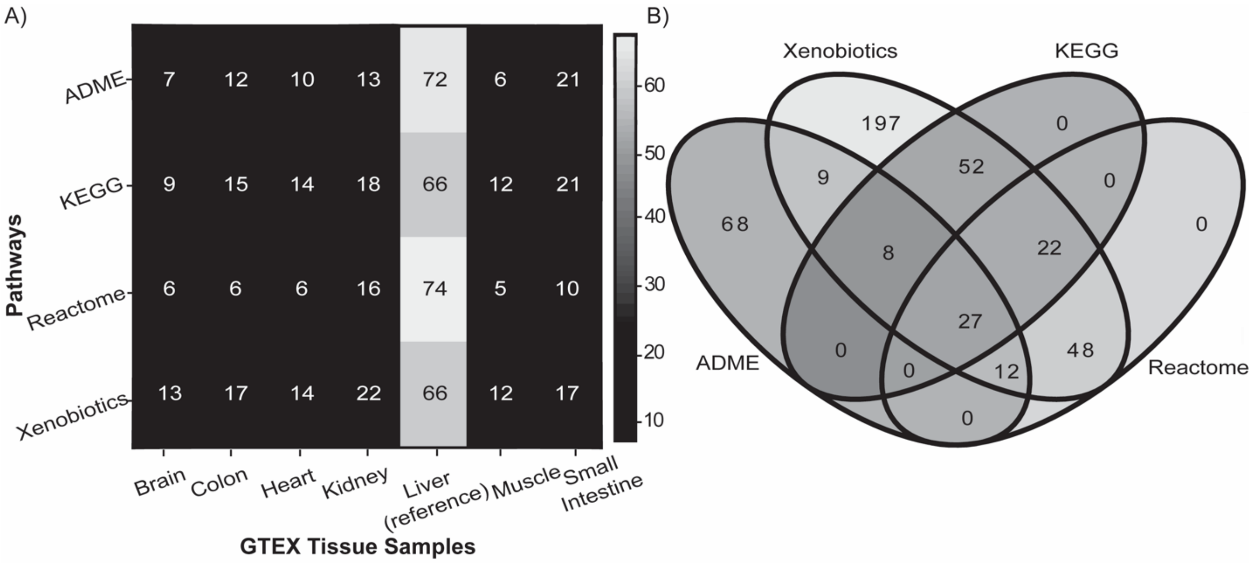

*Figure 3: Assessment of the effect of ADME gene sets obtained from various databases on the PBSM-based similarity score. A) Heatmap for median similarity scores for the different ADME pathway databases on the y-axis indicating the differences in the similarity scores between the different organ types on the x-axis. B) Venn diagram showing the gene distribution for ADME pathways from different databases.*

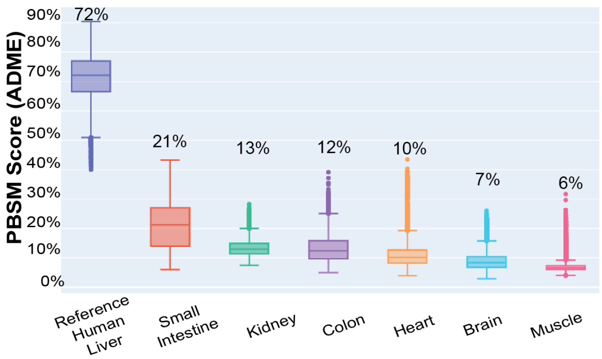

*Figure S2: PBSM-based similarity score distribution for Tissue samples from GTEX. Median similarity scores for GTEX tissue samples using the extended ADME geneset for the Reference Human Liver Tissue samples [Table S1].*

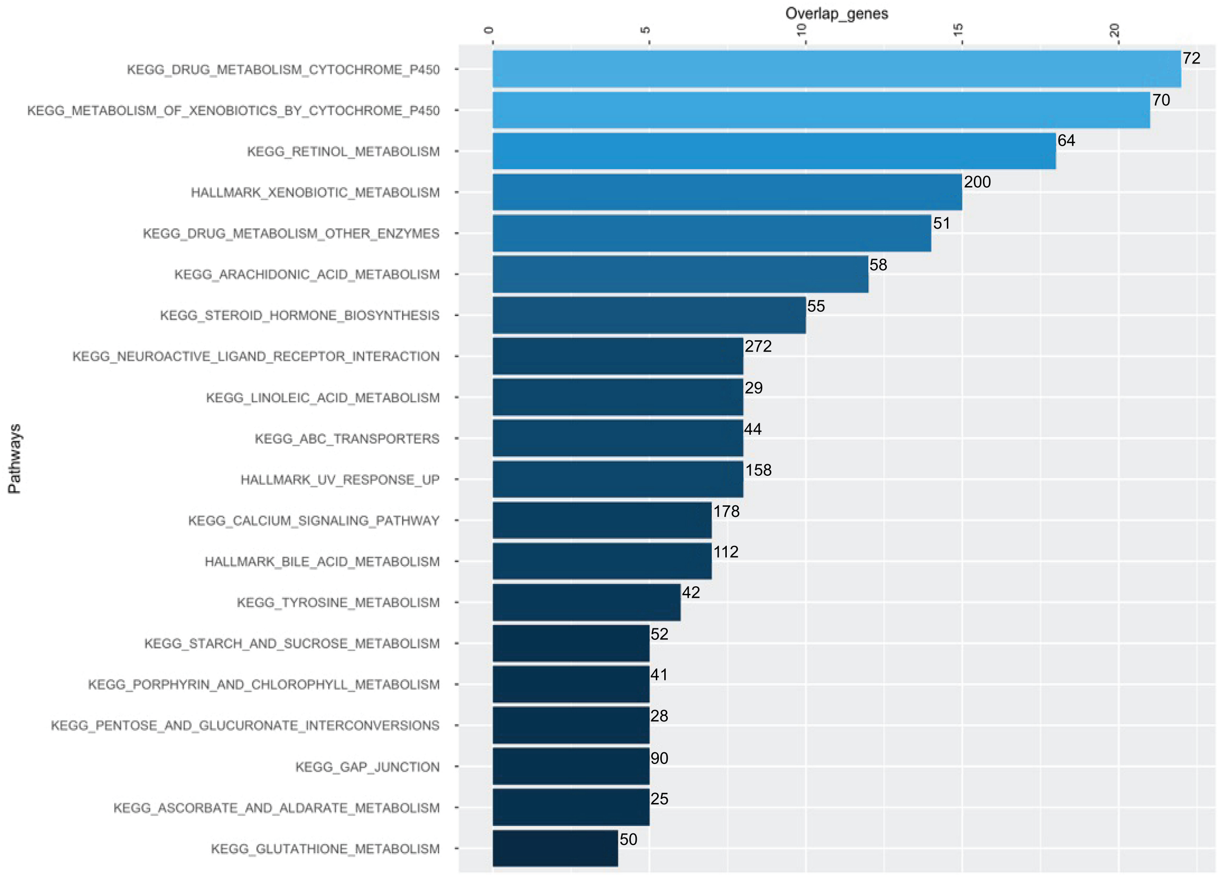

*Figure S3: Overlap analysis showing the shared genes between the extended ADME gene set and the top 20 pathways as defined by databases using the GSEA investigate feature [9][10].*

The extended ADME gene set was obtained from the [8] and defined in the Table S1. The PBSM score distribution for the extended ADME geneset defined in Table S1 can be seen in Figure S2. This figure shows Human small intestine and kidney tissue samples have the highest levels of similarity.

Figure S3 above shows the top 20 pathways described in various databases which were obtained from the overlap analysis using the GSEA investigate feature. The key pathways obtained from this analysis include those defined for the drug and xenobiotic metabolism.

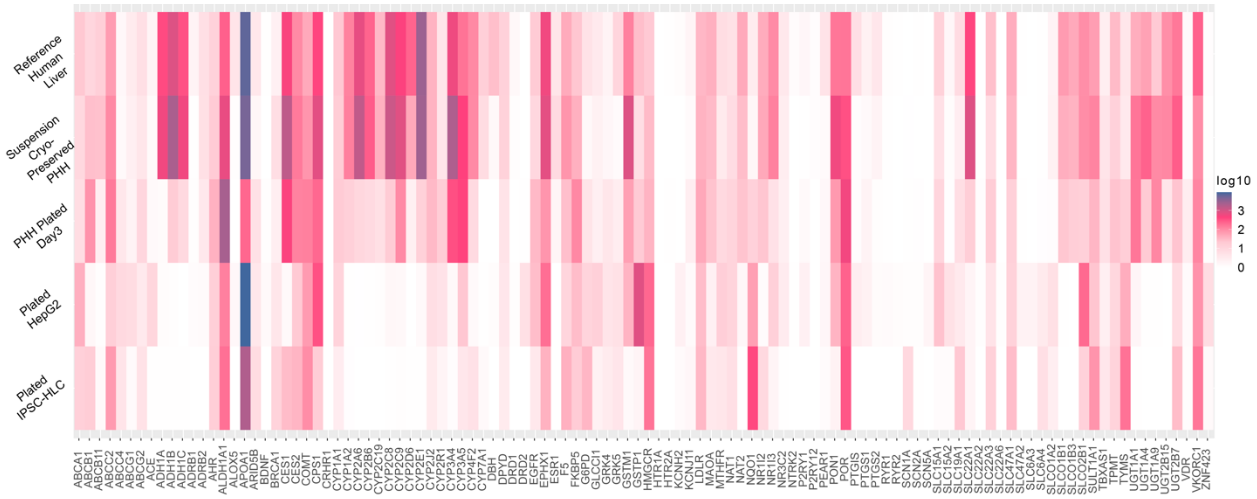

*Figure S4: Heatmap for the log10 Gene Expression using the genes defined in Table S3 for Figure 4a.*

The heatmap for the gene expression data in Figure S4 aligns with the results in Figure 4a. Figure 4a compares the PBSM similarity score of four *in-vitro* models to Reference Human Liver samples. Suspension cultures of human liver cells (PHH) were most similar to the reference, while Plated iPSC-HLC was the least similar, showing high and low gene expression in Figure S4 across the genes defined in Table S1 respectively.

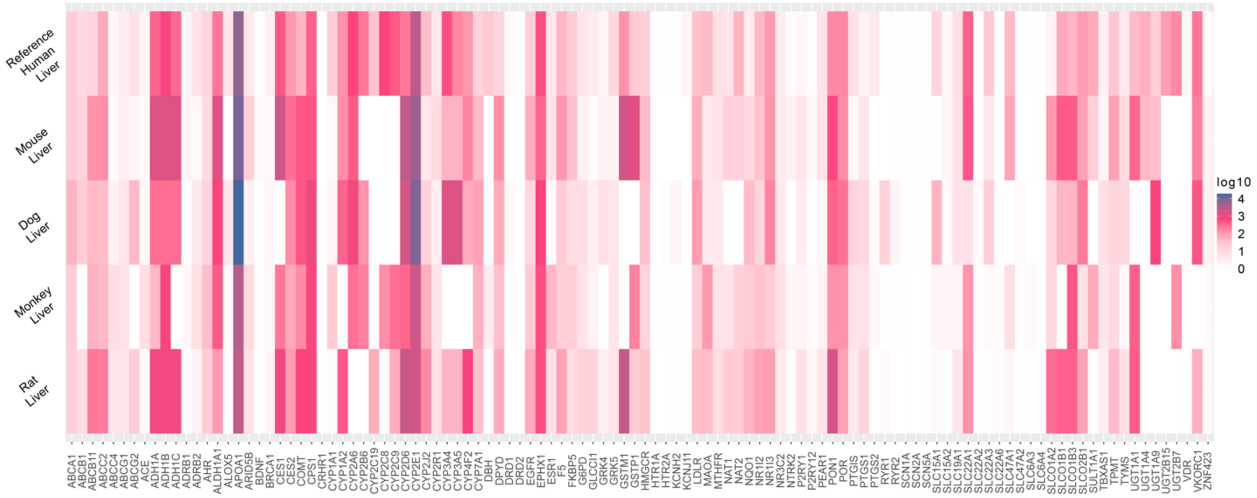

*Figure S5: Heatmap for the log10 Gene Expression using the genes defined in Table S1 for Figure 4b.*

The heatmap for the gene expression data in Figure S5 aligns with the results in Figure 4b. Figure 4b compares the PBSM similarity score of four *in-vivo* models to Reference Human Liver samples. Mouse liver tissue samples were most similar to the reference, showing high gene expression in Figure S5 across the genes defined in Table S1.

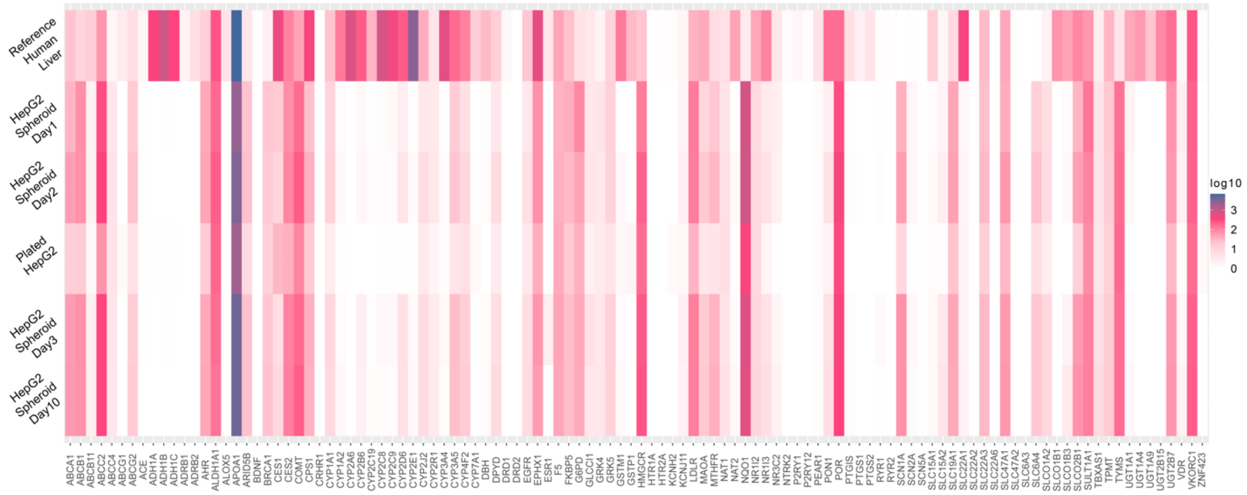

*Figure S6: Heatmap for the log 10 Gene Expression using the genes defined in Table S3 for Figure 4c.*

The heatmap for the gene expression data in Figure S6 aligns with the results in Figure 4c. Figure 4c compares the PBSM similarity score of the 3D and 2D models of HepG2 to Reference Human Liver samples. There were no significant differences between the HepG2 models, seen with no differences in the gene expression in Figure S6 across the genes defined in Table S1.

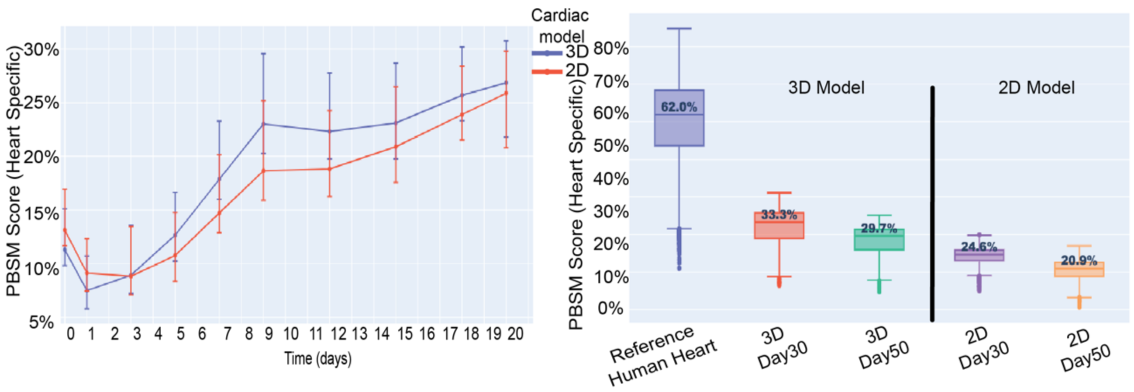

*Figure S7: PBSM-based similarity scores estimated for IPSc-derived cardiomyocytes A) Median similarity scores for both the 2D and 3D models during in vitro maturation for 20 days using custom cardiac-related genes for the Reference Human Ventricle Samples [Table S4]. B) Median similarity scores showing the effects of the long-term culture for the 2D and 3D cardiomyocytes until 50 days for the Reference Human Atrial Samples.*

Figure S7 consists of the pairwise comparison of the different iPSC-derived cardiomyocytes. The figure corresponds to Figure 5a and Figure 5b, where it compares the differences between the maturation for 2D and 3D models and the long-term effects for both models, respectively. The reference samples in Figure S7a are the Reference Human Ventricle Samples. For the Figure S7b, the reference samples are the Reference Human Atrial Samples.

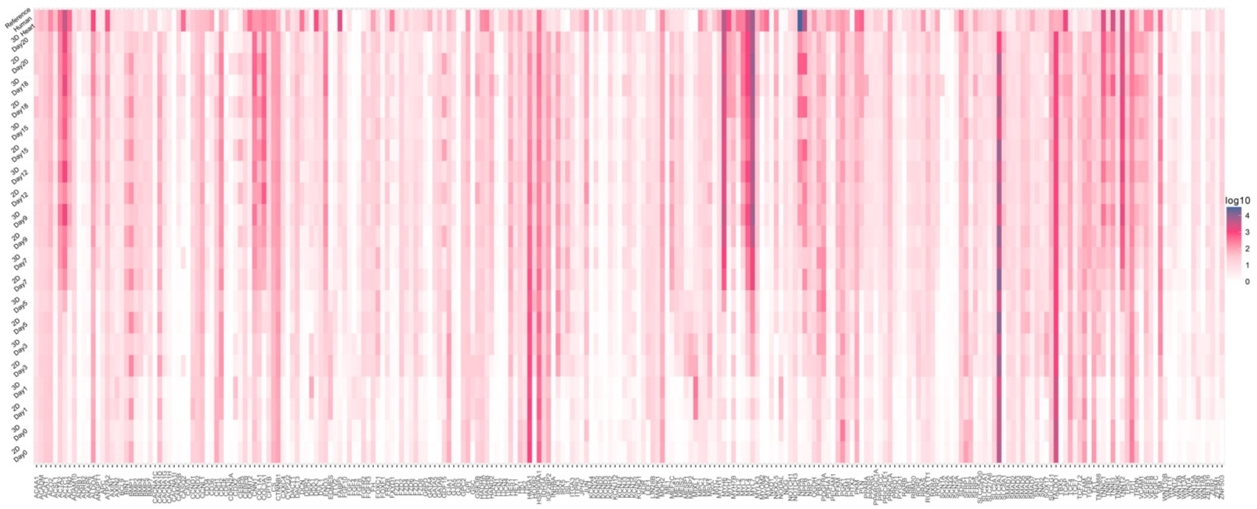

*Figure S8: Heatmap for the log10 Gene Expression for the Cardiac Maturation genes [11] defined in Table S4 for Figure 5a using the Reference Human Atrial Samples.*

The heatmap for the gene expression data in Figure S8 aligns with the results in Figure 5a. Figure 5a compares the PBSM similarity score of the 2D and 3D models of iPSC-derived cardiomyocytes to Reference Human Atrial Samples. Day 9 gene expression for 3D models was much higher than the corresponding 2D model when compared to the reference, corroborated by the differences in gene expression seen in Figure S8.

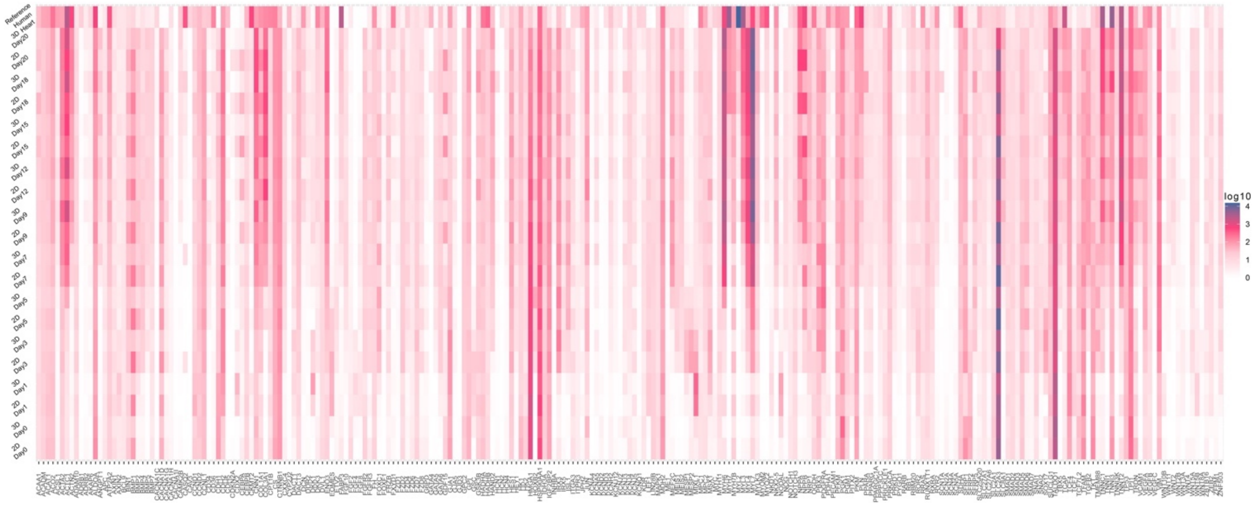

*Figure S9: Heatmap for the log10 Gene Expression for the Cardiac Maturation genes [11] defined in Table S4 for Figure S6a using the Reference Human Ventricle Samples.*

The heatmap for the gene expression data in Figure S9 aligns with the results in Figure S6a. Figure S6a compares the PBSM similarity score of the 2D and 3D models of iPSC-derived cardiomyocytes to Reference Human Ventricle Samples. Day 9 gene expression for 3D models was much higher than the corresponding 2D model when compared to the reference, corroborated by the gene expression in Figure S9.

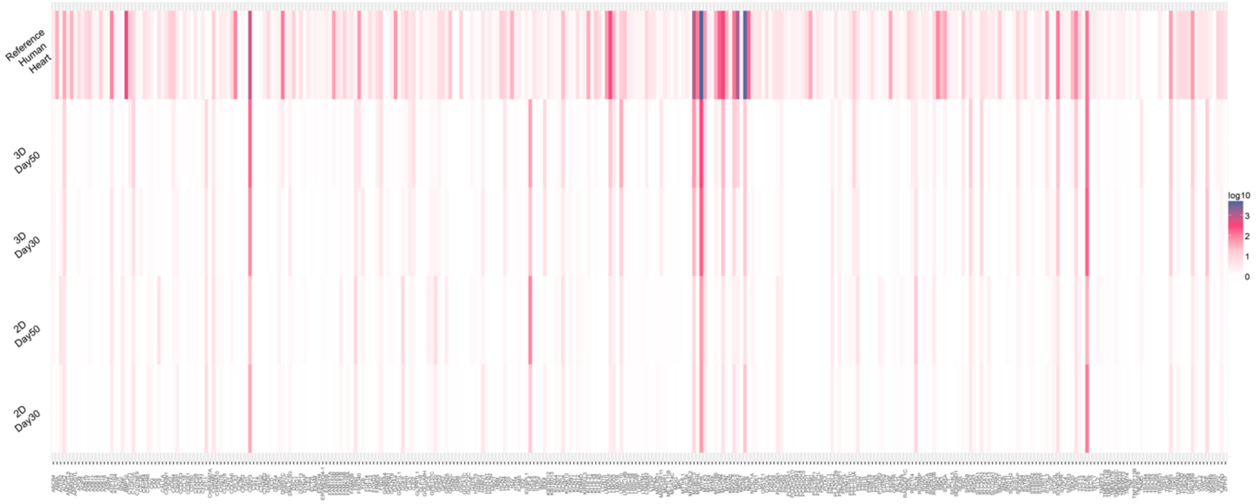

*Figure S10: Heatmap for the log10 Gene Expression for the genes defined specifically for Heart in the Human Protein Atlas (HPA)[12] for Figure S 6b using the Reference Human Atrial Samples.*

The heatmap for the gene expression data in Figure S10 aligns with the results in Figure S6b. Figure S6b compares the PBSM similarity score of the 2D and 3D models of iPSC-derived cardiomyocytes to Reference Human Atrial Samples. The 3D model maintains higher similarity for Day50 than the corresponding 2D model when compared to the reference, corroborated by the gene expression in Figure S10.

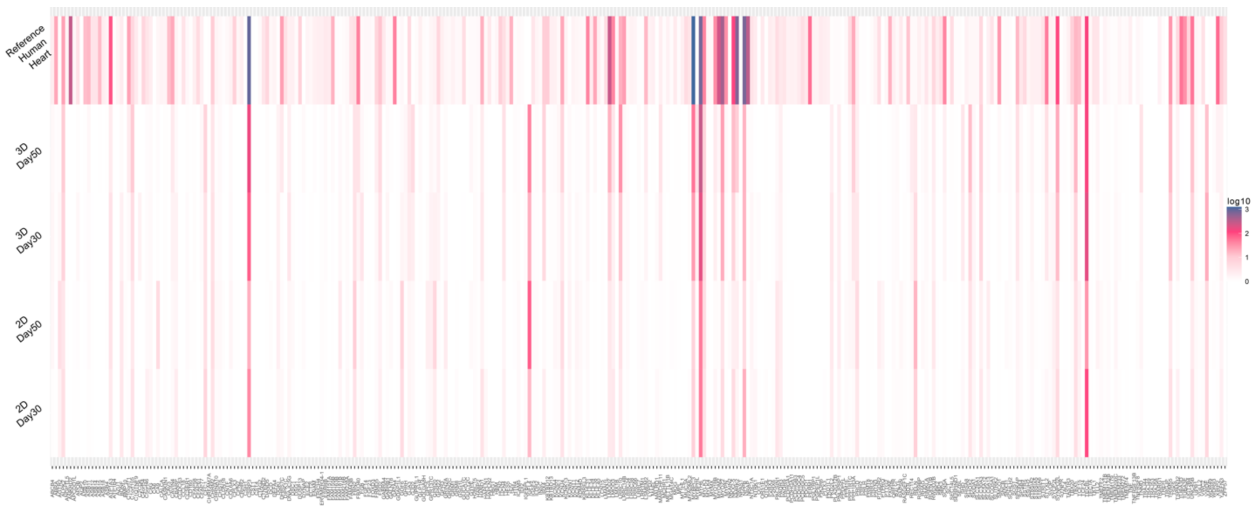

*Figure S11: Heatmap for the log10 Gene Expression for the genes defined specifically for Heart in the Human Protein Atlas (HPA)[12] for Figure 5b using the Reference Human Ventricle Samples.*

The heatmap for the gene expression data in Figure S11 aligns with the results in Figure 5b. Figure 5b compares the PBSM similarity score of the 2D and 3D models of iPSC-derived cardiomyocytes to Reference Human Ventricle Samples. The 3D model maintains higher similarity for Day50 than the corresponding 2D model when compared to the reference, corroborated by the gene expression in Figure S11.

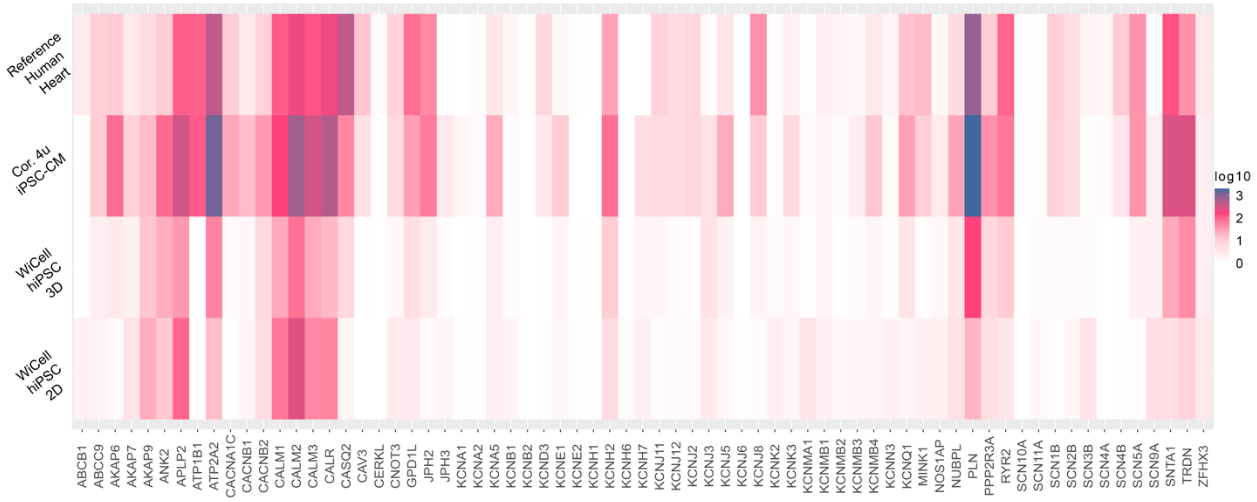

*Figure S12: Heatmap for the log10 Gene Expression for the genes affecting QT prolongation defined in the S.Table5 for Figure 5c using the Reference Human Ventricle Samples.*

The heatmap for the gene expression data in Figure S12 aligns with the results in Figure 5c. Figure 5c compares the PBSM similarity score of the different iPSC-derived cardiomyocytes models to Reference Human Ventricle Samples for their ability to predict drug-induced QT prolongation. The Cor4u. iPSC-CM were most similar to the reference, showing high gene expression in Figure S12 across the genes defined in Table S5.

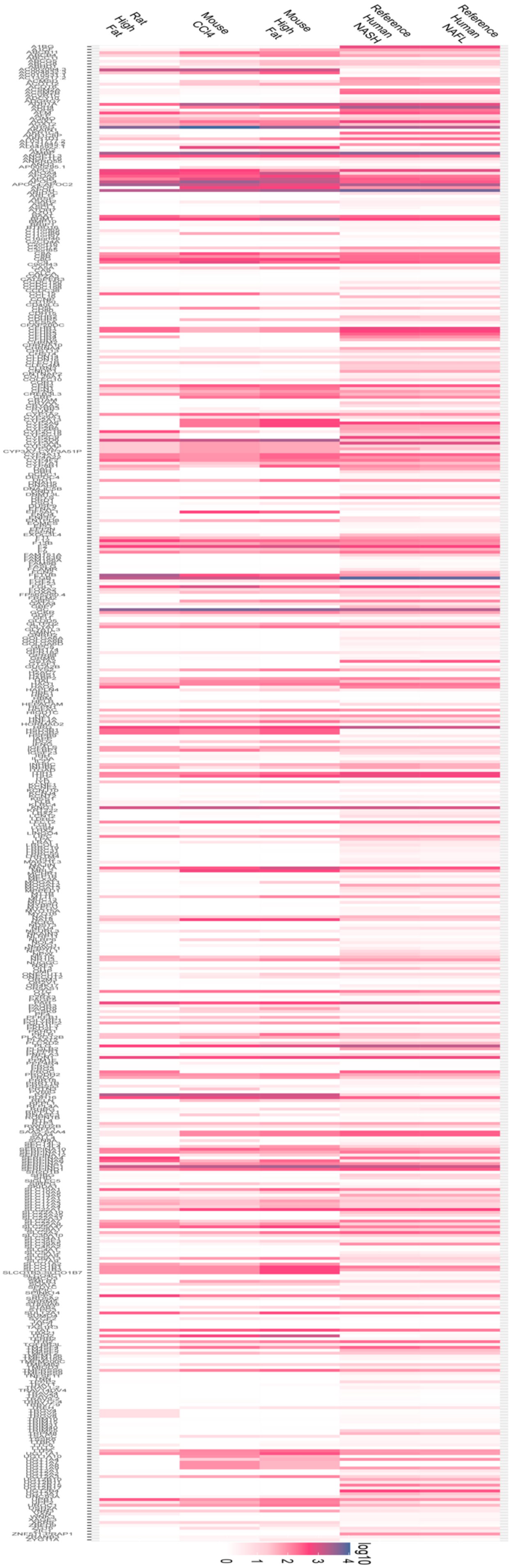
Rat High Fat
Mouse CCl4
Mouse High Fat
Reference Human NASH
Reference Human NAFL
log10
0
1
2
3
4

*Figure S13: Heatmap for the log10 Gene Expression for the genes defined specifically for the Liver in the HPA[12] for Figure 6a using the Reference Human NAFL samples.*

The heatmap for the gene expression data in Figure S13 aligns with the results in Figure 6a. Figure 6a compares the PBSM similarity score of the different in-vivo diseased models to Reference Human NAFL Samples. None of the in-vivo diseased models showed significant differences from the references for the liver-specific genes, showing similar low gene expression in Figure S13.

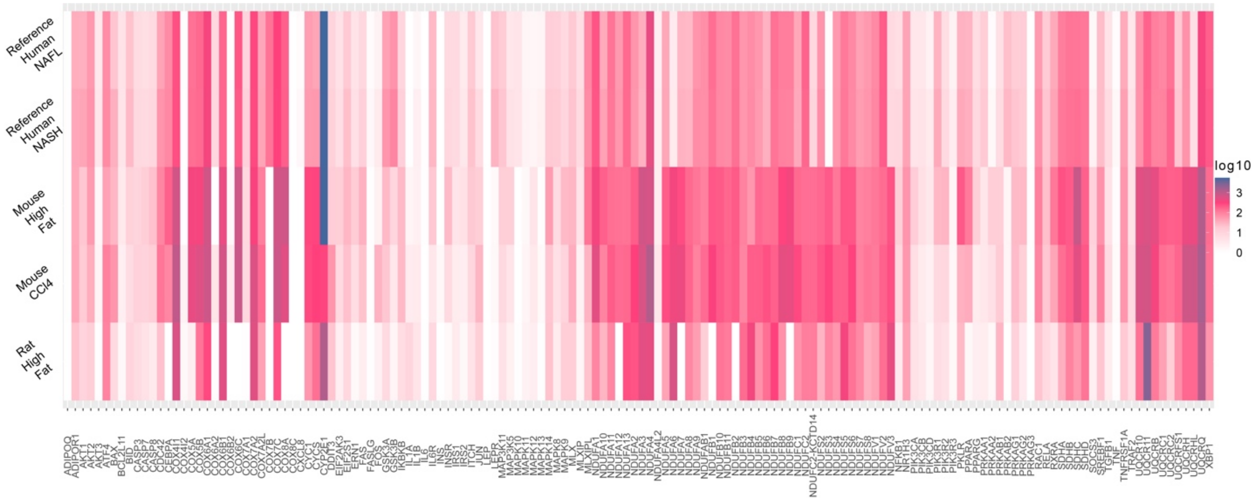

*Figure S14: Heatmap for the log10 Gene Expression for the genes defined in S.Table6 for Figure 6b using the Reference Human NAFL samples.*

The heatmap for the gene expression data in Figure S14 aligns with the results in Figure 6b. Figure 6b compares the PBSM similarity score of the different in-vivo diseased models to Reference Human NAFL Samples. The Mouse High Fat diseased model was the most similar to the reference, showing high gene expression in Figure S14 across the genes defined in Table S6.

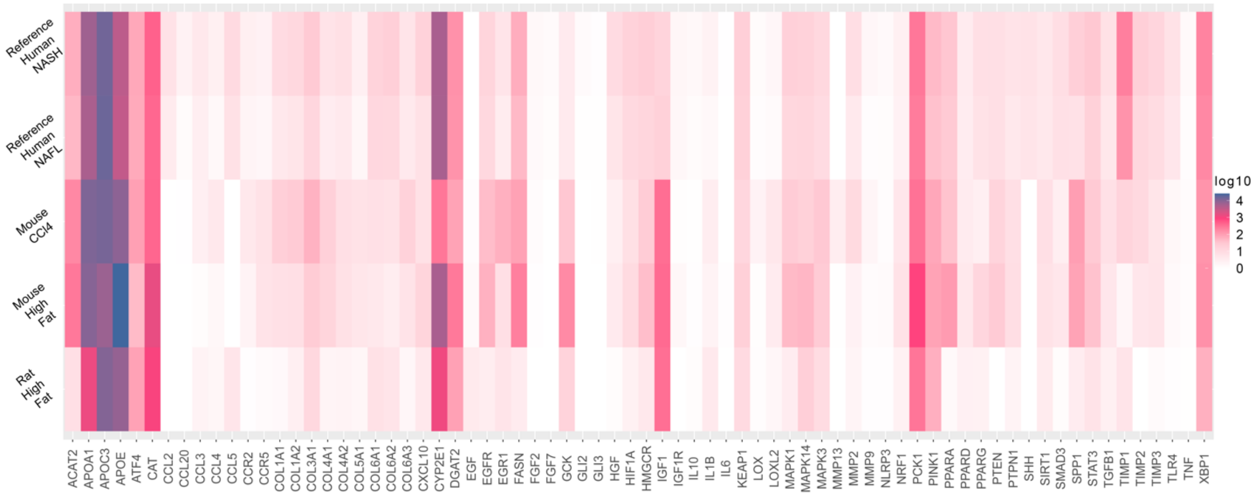

*Figure S15: Heatmap for the log10 Gene Expression for the genes defined in S.Table7 for Figure 6c using the Reference Human NASH samples.*

The heatmap for the gene expression data in Figure S15 aligns with the results in Figure 6c. Figure 6c compares the PBSM similarity score of the different in-vivo diseased models to Reference Human NASH Samples. The Mouse model injected with CCl4 was the most similar to the reference, showing high gene expression in Figure S15 across the genes defined in the Table S7.

**References:**

1. Andrews, Simon. "FastQC: a quality control tool for high throughput sequence data." (2010).
2. Chen S, Zhou Y, Chen Y, et al. fastp: an ultra-fast all-in-one FASTQ preprocessor. Bioinformatics. 2018; 34(17)' i884-i890. doi: 10.1093/bioinformatics/bty560.
3. Patro R, Duggal G, Love MI, et al. Salmon provides fast and bias-aware quantification of transcript expression. Nat Methods. 2017; 14(4); 417-419. doi: 10.1038/nmeth.4197.
4. Durinck S, Spellman PT, Birney E, et al. Mapping identifiers for the integration of genomic datasets with the R/Bioconductor package biomaRt. Nat Protoc. 2009;4(8):1184-91. doi: 10.1038/nprot.2009.97.

5. "Human Gene Set." KEGG_DRUG_METABOLISM_CYTOCHROME_P450, www.gsea-msigdb.org/gsea/msigdb/cards/KEGG_DRUG_METABOLISM_CYTOCHROME_P450. Accessed 02 Jun. 2022.
6. KEGG PATHWAY: Map00983, www.genome.jp/entry/pathway+map00983. Accessed 02 Apr. 2024.
7. Bijay, et al. "Reactome: Drug ADME." Reactome Pathway Database, reactome.org/content/detail/R-HSA-9748784. Accessed 02 Jun. 2022.
8. Han SM, Park J, Lee JH, et al. Targeted Next-Generation Sequencing for Comprehensive Genetic Profiling of Pharmacogenes. Clin Pharmacol Ther. 2017; 101(3); 396-405. doi: 10.1002/cpt.532.
9. Liberzon A, Birger C, Thorvaldsdóttir H, et al. The Molecular Signatures Database (MSigDB) hallmark gene set collection. Cell Syst. 2015; 1(6); 417-425. doi: 10.1016/j.cels.2015.12.004.
10. Subramanian A, Tamayo P, Mootha VK, et al. Gene set enrichment analysis: a knowledge-based approach for interpreting genome-wide expression profiles. Proc Natl Acad Sci USA. 2005; 102(43); 15545-50. doi: 10.1073/pnas.0506580102.
11. Branco MA, Cotovio JP, Rodrigues CAV, et al. Transcriptomic analysis of 3D Cardiac Differentiation of Human Induced Pluripotent Stem Cells Reveals Faster Cardiomyocyte Maturation Compared to 2D Culture. Sci Rep. 2019; 9(1); 9229. doi: 10.1038/s41598-019-45047-9.

12. Uhlén M., Fagerberg L., Björn M., et al. Tissue-Based Map of the Human Proteome. Science 347 2015; 6220; 1260419. doi: 10.1126/science.1260419. https://www.proteinatlas.org/.
13. KEGG PATHWAY: Hsa04932, www.genome.jp/entry/pathway+hsa04932. Accessed 10 Apr. 2023.
